# Circuit to target approach defines an autocrine myofibroblast loop that drives cardiac fibrosis

**DOI:** 10.1101/2023.01.01.522422

**Authors:** Shoval Miyara, Miri Adler, Elad Bassat, Yalin Divinsky, Kfir B. Umansky, Jacob Elkahal, Alexander Genzelinakh, David Kain, Daria Lendengolts, Tali Shalit, Michael Gershovits, Avraham Shakked, Lingling Zhang, Jingkui Wang, Danielle M. Kimchi, Andrea Baehr, Rachel Sarig, Christian Kupatt, Elly M. Tanaka, Ruslan Medzhitov, Avi Mayo, Uri Alon, Eldad Tzahor

## Abstract

Fibrosis is a broad pathology of excessive scarring with substantial medical implications. The fibrotic scar is produced by myofibroblasts that interact with macrophages. Fibrosis is a complex process involving thousands of factors, therefore, to better understand fibrosis and develop new therapeutic approaches, it is necessary to simplify and clarify the underlying concepts. Recently, we described a mathematical model for a macrophage-myofibroblast cell circuit, predicting two types of fibrosis - hot fibrosis with abundant macrophages and myofibroblasts, and cold fibrosis dominated by myofibroblasts alone. To test these concepts and intervention strategies in a medically relevant system, we use a widely studied *in-vivo* injury model for fibrosis, myocardial infarction (MI). We show that cold fibrosis is the final outcome of MI in both mice and pigs and demonstrate that fibrosis can shift toward healing in regenerative settings. MI begind with an increase of myofibroblasts and macrophages, followed by macrophage decline leading to persistent cold fibrosis (only myofibroblasts). During this process, fibroblasts, unlike macrophages, acquire distinct fate changes. Using mathematical modeling we predict that targeting of the autocrine signal for myofibroblast division could block cold fibrosis. We identify TIMP1 as an autocrine cardiac myofibroblast growth factor *in-vitro*. Treatment of adult mice after MI with anti-TIMP1 antibodies reduces fibrosis *in-vivo*. This study shows the utility of the concepts of hot and cold fibrosis and the feasibility of our circuit-to-target approach to reduce fibrosis after acute cardiac injury by inhibiting the myofibroblast autocrine loop.

## INTRODUCTION

Fibrosis is a common pathology in which excessive scarring replaces healthy tissue (Henderson, Rieder, & Wynn, 2020; T. Wynn, 2008). Fibrosis gradually leads to organ failure in many tissues, including heart (Davis & Molkentin, 2014), kidney (Eddy, 2014), liver (S. L. Friedman, 2008), muscle (Mann et al., 2011), and lung (Adams et al., 2020; Noble, Barkauskas, & Jiang, 2012). Preventing or reducing fibrosis is a major unmet medical need (Henderson et al., 2020; T. Wynn, 2008). Fibrosis is a complex biological process due to the interaction of multiple cell types and signaling molecules (Duffield, Lupher, Thannickal, & Wynn, 2013; T. A. Wynn & Vannella, 2016). The structural component of the pathology, the scar, is composed of extracellular matrix (ECM) proteins deposited mostly by activated fibroblasts, called myofibroblasts (Klingberg, Hinz, & White, 2013). Myofibroblasts interact with many cell types including monocyte-derived macrophages ((Buechler, Fu, & Turley, 2021; Pakshir & Hinz, 2018), which are recruited in large numbers into the site of injury and support the fibrotic process (Lech & Anders, 2013; Pakshir & Hinz, 2018; Tzahor & Dimmeler, 2022; T. A. Wynn & Vannella, 2016) .

Recently we developed a mathematical model of fibrosis, focusing on a circuit of growth factor exchange between myofibroblasts and macrophages (Adler et al., 2020; Zhou et al., 2018). The model suggests three possible outcomes - healing, hot fibrosis, or cold fibrosis. Hot fibrosis is characterized by both myofibroblasts and macrophages which support each other’s growth. Cold fibrosis, in contrast, consists of myofibroblasts without activated macrophages. Healing is accompanied by collapse of the two cell populations and return to baseline (i.e., tissue homeostasis). The model predicts that during cold fibrosis, myofibroblast growth is driven by an autocrine growth factor loop (Adler et al., 2020). The model also predicts that suitable modulation of the growth factor milieu can provide transitions between healing and fibrosis states. However, the relevance of hot and cold fibrosis and their therapeutic modulation has not been tested in a clinically relevant system.

We sought to test these concepts and the ability to manipulate them for therapy using a widely studied *in-vivo* model - fibrosis after myocardial infarction (MI). MI occurs when a blood vessel that feeds the heart is blocked, causing ischemia and cardiomyocyte cell death (Frangogiannis, 2018; Reed, Rossi, & Cannon, 2017). Subsequently, immune cells rapidly invade the injured zone leading to the local expansion and activation of myofibroblasts. This results in changes in the microenvironment dictated by the release of various cytokines, growth factors and matrix proteins. The injured myocardium is replaced by fibrotic scar over a period of several weeks, a process called cardiac remodeling, that gradually leads to reduced heart function, morbidity and mortality (Heymans et al., 2015; Hinderer & Schenke-Layland, 2019; Stuart, Jesus, Lindsey, & Ripplinger, 2016).

Cardiac fibrosis is an inevitable outcome in adult mammals upon most cardiac injuries or stresses, but is absent in neonatal mice and pigs due to the regenerative potential of the young heart (Haubner et al., 2016; Porrello et al., 2011; Tzahor & Poss, 2017; Ye et al., 2018; Zhu et al., 2018). Recent advances in animal models of cardiac injuries suggest that fibrosis can be reduced by certain interventions (Aghajanian et al., 2019; Aharonov et al., 2020; Alexanian et al., 2021; Bassat et al., 2017; D’Uva et al., 2015; Gabisonia et al., 2019; Leach et al., 2017; Nakada et al., 2017; Sadek & Olson, 2020; Vagnozzi et al., 2020) - changing a long-held notion in the field that fibrosis (or scarring) is irreversible. These interventions have yet to be translated to the clinic.

Here we use bulk, single-cell and spatial transcriptomics in mice and pigs to show that fibrosis after MI is a dynamic process that ends in cold fibrosis, dominated by myofibroblasts without activated macrophages. We use the mathematical circuit model to identify a target to reduce cold fibrosis - the myofibroblast autocrine loop. We then identify the growth factors that contribute to this autocrine loop using transcriptomic analysis and *in-vitro* experiments. We show that inhibiting one of these factors, TIMP1, by means of neutralizing antibodies reduces fibrosis after MI in mice. This study highlights the concept of cold fibrosis and the ability of a circuit-to-target approach to inhibit the myofibroblast autocrine loop and to reduce fibrosis after acute cardiac injury.

## RESULTS

### Acute cardiac injury results in cold fibrosis

Our starting point for discovering a target for MI-induced fibrosis is a quantitative theory of inflammation and fibrosis (Adler et al., 2020) (**Fig. 1A**). This theory is based on a dynamical model for the growth and interaction of myofibroblasts and macrophages determined from *in-vitro* co-cultures (Zhou et al., 2018). The state of the tissue is described in a phase portrait whose axes are the abundance of myofibroblasts and macrophages (**Fig. 1A**). Arrows mark the dynamics of the cells over time. The system shows three possible outcomes for injury. All three start by macrophage infiltration, accompanied by fibroblast activation and proliferation leading to an increase in both cell types. In healing (i.e., a regenerative setting) both populations eventually shrink back to pre-injury levels, providing a trajectory that rises and then falls in the phase portrait (**Fig. 1A**, blue line). The other two outcomes are fibrotic states. In hot fibrosis myofibroblasts and macrophages support each other’s growth by a reciprocal exchange of growth factors, providing a trajectory that rises to a fixed point with high levels of both myofibroblasts and macrophages (**Fig. 1A**). The other possible fibrotic outcome is cold fibrosis. Both macrophages and myofibroblasts initially rise, but then macrophage numbers return to baseline while myofibroblasts persist. Cold fibrosis is thus a state of abundant myofibroblasts that support their own growth, with low macrophages similar to the pre-injury state (**Fig. 1A**).

**Figure 1:**
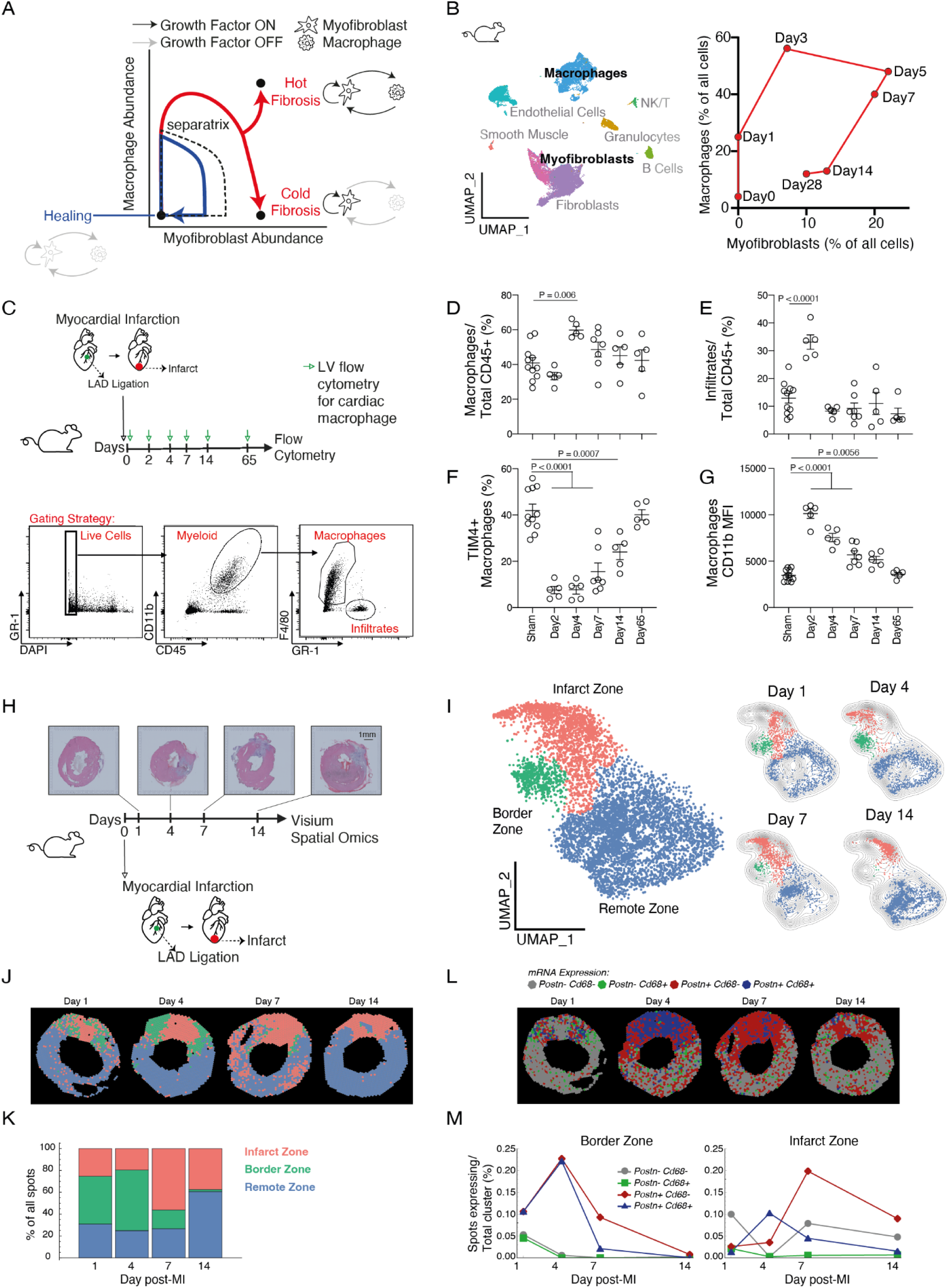
Acute myocardial infarction results in cold fibrosis. **(A)** Mathematical model of the myofibroblast-macrophage circuit, (adapted from (Adler et al., 2020), predicts three possible outcomes - hot fibrosis, cold fibrosis and healing based on the abundance of these two populations (axes). The basin of attraction for the healing state is bounded by a separatrix. The scheme also denotes the growth factor signaling as either ON (black lines) or OFF (grey lines) at each specific fixed point. **(B)** Single-cell RNA-sequencing (scRNAseq) dynamic analysis of total macrophage and myofibroblast populations (% of total interstitial cells) in an adult mouse heart following-MI data obtained from Forte et al., (2020). These data are further described in fig. S1A-B. UMAP of clusters, corresponding to: Smooth Muscle Cells, Macrophages, Fibroblasts, Myofibroblasts, Endothelial Cells, Granulocytes, NK/T cells and B cells. **(C)** Top panel- experimental scheme whereby adult mice hearts underwent MI by permanent left anterior descending artery (LAD) and further processed by flow cytometry. Left ventricles (LV) of MI or Sham operated mice were collected at 6 different timepoints following injury at days: 0/Sham (n = 11), 2 (n = 5), 4 (n = 5), 7 (n = 7), 14 (n = 5) and 65 (n = 5). Bottom panel- representative flow cytometry plots of the gating scheme used to identify **(D)** total cardiac macrophages (out of total immune cells- CD45^+^), **(E)** Infiltrating cells (monocytes and neutrophils out of total immune cells-CD45^+^), **(F)** Resident macrophages (TIM4^+^ macrophages out of total macrophages) and **(G)** cardiac macrophage activation, measured by CD11b median fluorescence intensity (MFI) (Orrskog et al., 2012). Data presented as mean± SEM **(H)** Experimental design of adult mice hearts underwent MI and processed by Visium spatial transcriptomics analysis at 4 different time points following injury [at days: 1 (n = 1), 4 (n = 1), 7 (n = 1) and 14 (n = 1)]; Top panel: representative Hematoxylin and Eosin (H&E) stained section. Scale bar, 1 mm. **(I)** Left panel- uniform manifold approximation and projection (UMAP) plot of total spots from all timepoints combined day1 (n = 1516), day 4 (n = 1596), day 7 (n = 1510), day 14 (n = 2117) . Spots were annotated as 3 distinct gene-expression clusters [Infarct Zone (orange), Border Zone (green), Remote (Healthy) Zone (blue)]. Right panel- UMAP plots of the 3 clusters according to time following MI with density contour based on smooth kernel distribution. **(J)** Clusters (Infarct, border and remote zones) represented by their spatial coordinates per timepoint. **(K)** Cluster distribution (% of total spots per timepoint). **(L)** Macrophage and myofibroblast spatio-temporal distribution after MI. Spots were defined as macrophage (*Cd68*^+^; *Postn*^-^; green), myofibroblast (*Cd68*^-^; *Postn*^+^; red), macrophage+myofibroblast (*Cd68*^+^; *Postn*^+^; blue) or spots without macrophages or myofibroblasts (*Cd68*^-^; *Postn*^-^; grey). Positive spots (either *Cd68*, *Postn* or both) were defined by expression, of each gene, higher than median over all slices. **(M)** Quantification of macrophage-myofibroblast spot distribution presented in (L) and divided per zone (Left panel- border zone or right panel- infarct zone). Results are presented as % of spots out of the total spots per spatial cluster. For all spatial spot distribution plots (J and L) each slice was rotated so that the infarct zone median (defined by infarct zone spots) is positioned upwards.

Whether hot or cold fibrosis occurs depends on the tissue parameters in the model and the type of injury. For example, a strong paracrine support of macrophages by myofibroblasts leads to hot fibrosis, whereas a weaker paracrine support is predicted to result in cold fibrosis (Adler et al., 2020). We hypothesize that the pathology and therapeutic options for fibrosis should differ in a setting of hot or cold fibrosis.

To explore hot and cold fibrosis in a clinically relevant setting, we use acute-MI as a model system in which fibrosis gradually develops after injury (Stuart et al., 2016). We first explored the dynamics of macrophage and myofibroblast populations in mice after MI. To do so, we analyzed single cell RNA-sequencing data (Forte et al., 2020) and quantified the abundance of macrophages and myofibroblasts at seven time points after MI. Macrophages and myofibroblasts first rise and peak at days 3-7. Then macrophages decrease leaving persistent myofibroblasts at days 14 and 28 (**Fig. 1B, fig. S1A-B**) (Fu et al., 2018). Day 28 is considered a mature scar. This trajectory is consistent with cold fibrosis.

To further test whether cold fibrosis is the endpoint of MI, we asked whether macrophages return to baseline. To analyze macrophage dynamics, we performed MI on adult mice (3-month old), harvested cells from the injured left ventricle at different timepoints, and analyzed them by flow cytometry using cardiac macrophage markers (**Fig. 1C**). The macrophage fraction relative to all immune cells (CD45^+^) increased and then returned to baseline levels, as did the fraction of infiltrating monocytes and neutrophils (**Fig. 1D-E**) and the fraction of activated macrophages (**Fig. 1G**). In contrast, tissue-resident macrophages which are required for cardiac regeneration and repair (Aurora et al., 2014; Dick et al., 2019; Lavine et al., 2014) were depleted rapidly but gradually restored 65 days post MI to baseline levels (**Fig. 1F**). The cardiac macrophage population thus returns to baseline in the aftermath of MI, consistent with a cold fibrosis trajectory.

To explore the spatial organization of cold fibrosis, we performed Visium transcriptomic analysis on 10μm thin tissue heart sections of 3-month old mice post-MI. Visium allows spatially resolved gene expression by performing mRNA-sequencing on 55μm spots arrayed across each section (**Fig. 1H**). We find three gene-expression clusters by unbiased clustering (**Fig. 1I, methods**). The spatial position of these clusters indicates that they correspond to the infarct zone (with scar components such as: *Col1a1*, *Col3a1, Fn1*), remote zone (with functional cardiomyocyte marker genes such as: *Myh6, Myl2*) and a defined border zone population between them (with marker genes such as *Nppa*, *Ankrd1* and *Clu)* (Calcagno et al., 2022; Duijvenboden et al., 2019; Yamada et al., 2022) (**fig. S1D-E, Table S1**). The border zone population peaks at day 4, and then shrinks in size (**Fig. 1I-K**).

Marker genes showed that macrophages (*Cd68*) and myofibroblasts (*Postn*) were upregulated in the border (day 4) and infarct (day 7) zones and then the macrophage population decreased, whereas myofibroblasts persisted in the infarct zone (**Fig.1L-M**). Thus, the dynamic response to MI injury includes two distinct phases - an inflammatory phase with macrophages and a distinct border zone at days 1 to 7 post MI, followed by cold fibrosis dominated by myofibroblasts in the infarct zone and reduced border zone.

### Cold fibrosis is conserved in pigs

To determine whether cold fibrosis following MI is conserved across species we used a clinically relevant porcine model of cardiac injury. Pigs underwent a 60 minutes ischemia-reperfusion injury by the inflation of a balloon within the left anterior descending (LAD) coronary artery (Hinkel et al., 2015). Following reperfusion, pigs were treated, in an antegrade intracoronary trajectory, with saline solution (control) or with recombinant human Agrin protein (rhAgrin), a regenerative ECM protein we have previously shown to promote cardiac repair in mice and pigs (Baehr et al., 2020; Bassat et al., 2017). We obtained heart samples from healthy remote myocardium (Remote Zone) and the infarcted tissue (Infarct Zone) at days 3 and 28 following MI. We then performed bulk mRNA-sequencing and histological scar analysis (**Fig. 2A**).

**Figure 2:**
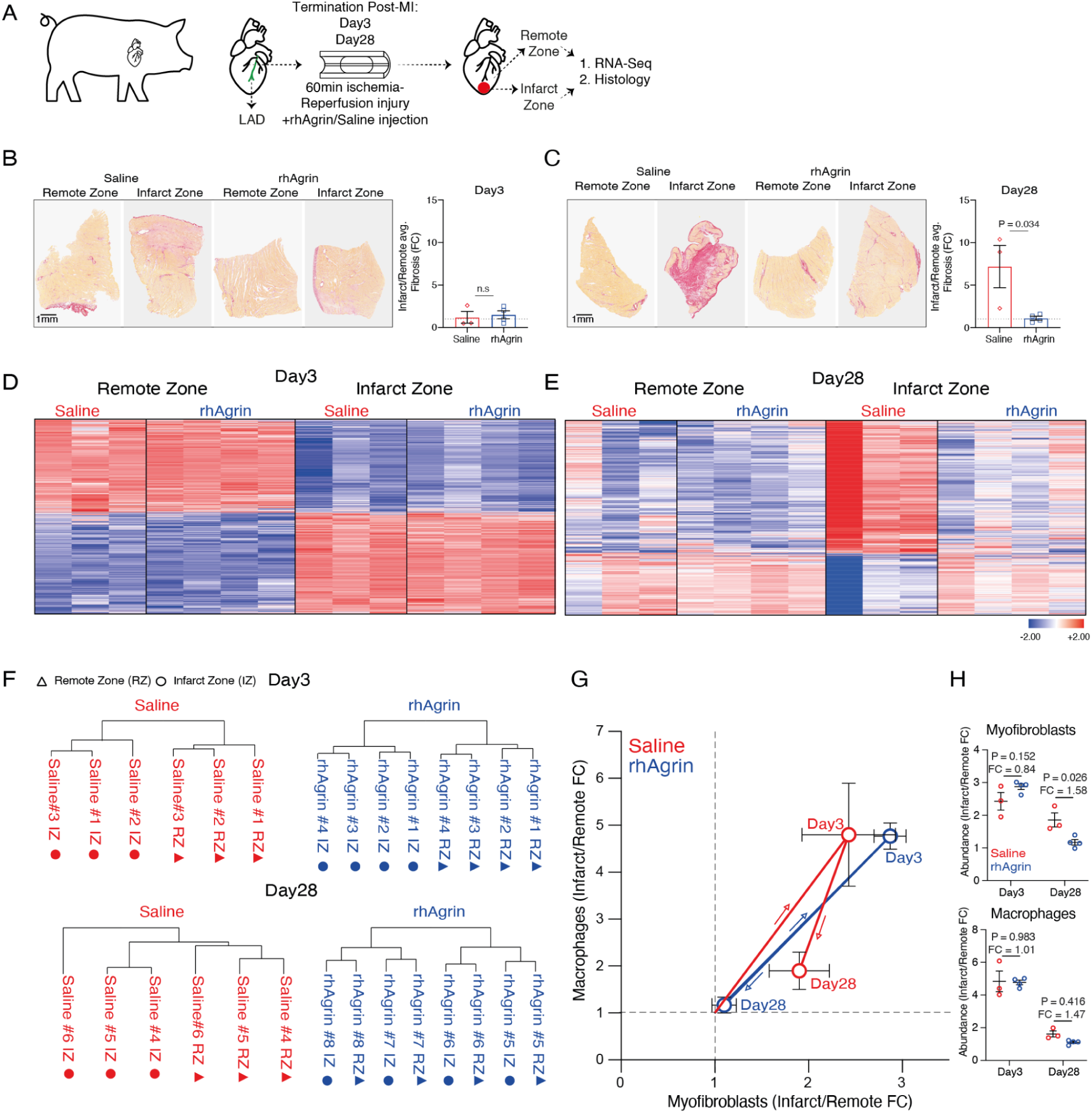
Cold fibrosis after MI is conserved in a clinically relevant porcine model. **(A)** Experimental design of pig MI experiment: adult (3 months old) pigs underwent MI by temporarily occluding their LAD using a balloon catheter (Methods). Following reperfusion (balloon deflation), pigs were immediately treated with recombinant human agrin (rhAgrin) or saline control, in an antegrade trajectory. Injured pig hearts were collected at either day 3 (n = 4 for rhAgrin; n = 3 for saline) or day 28 (n = 4 for rhAgrin; n = 3 for saline), and dissected to distinct tissue areas (Infarct and Remote zones). Remote and infarcted samples were subjected to bulk mRNA sequencing and histology for fibrosis assessment. **(B)** Representative picro-sirius red staining images are shown from day 3 and **(C)** day 28 post-MI. Fibrosis was quantified as the average % fold change (FC) between Infarct/Remote zone sections for each pig individually. Fibrosis (day 3 or 28) for Saline and rhAgrin hearts was measured using two-tailed unpaired Student t-test. Scale bars: 1 mm. striated line denotes FC = 1. n.s- non-significant difference. Heatmaps based on log2 transformed normalized counts for all (up and down regulated) differentially expressed genes (defined by |log2FoldChange| >= 1, p-adjusted value < 0.05 and max raw counts > 30) between remote and infarct zones for saline and rhAgrin treated hearts at day 3 (5961 genes) **(D)** and 28 (1079 genes) **(E)** post-MI. Rows represent genes and columns represent each biological sample and its spatial distribution according to infarct or remote zone. Data are represented as mean± SD. **(F)** Hierarchical clustering per condition [day (3 or 28) +treatment (agrin or saline)], based on the 1000 most variable genes. Triangles and circles represent remote and infarct zones, respectively. **(G)** Deconvolution of Bulk mRNA sequencing of pig hearts following MI of either rhAgrin (blue) or saline treated (red) samples. Macrophage and myofibroblast abundances were assessed by gene signatures as fold change(FC),between infarct and remote zones (Methods, Table S6). **(H)** Macrophage and myofibroblast abundances, based on deconvolution of bulk-mRNA sequencing (as in G). Treated samples were compared per time point (day 3 or 28), separately by two-tailed unpaired Student t-test. Results are represented as mean± SEM.

As expected, hearts treated with saline showed extensive picro-sirius red staining indicating severe fibrosis at day 28, whereas hearts treated with rhAgrin had less fibrosis, consistent with our previous findings (Baehr et al., 2020) (**Fig. 2B-C, fig. S2A-B**). Unlike saline treated hearts, rhAgrin treated infarct zone samples show a similar gene expression profile to their paired healthy remote zones at day 28 (**Fig. 2D-F, fig. S2D-I, Table S2-5**). We next used a deconvolution algorithm (**methods, fig. S2C, Table S6**) on the mRNA profiles to obtain the abundances of myofibroblasts and macrophages at both time points (**Fig. 2H-I**). As seen in mice after MI, saline treated pig hearts had increased macrophage abundance at day 3 which declined by day 28 in the infarct region, whereas myofibroblasts abundance remained high at 28 days indicative of cold fibrosis. In contrast, rhAgrin-treated hearts showed a healing trajectory in which both myofibroblast and macrophage abundance increased at day 3, and then returned to baseline levels at day 28, similar to the healing loop predicted by the model (**Fig. 1A**). We conclude that the natural dynamics of acute MI leads to cold fibrosis in both mice and pigs.

### In cold fibrosis macrophages return to a homeostatic state whereas fibroblasts acquire a profibrotic phenotype

We next asked whether cold fibrosis entails changes in cell states of macrophages and fibroblasts. Specifically, we analyzed gene expression states of the two populations and tracked their dynamics after MI using single-cell mRNA-sequencing (scRNAseq) data (Forte et al., 2020) from cardiac interstitial cells following MI (**Fig. 1B, fig. S1A-B**). We analyzed the gene expression of fibroblasts (including myofibroblasts) and macrophages at day 0 (i.e., pre-injury) and day 28 when fibrosis matures. Each cell population is a continuum in gene expression space, and therefore we defined the main functions of each cell type using Pareto task inference (ParTI) (**Fig. 3A**) (Hart et al., 2015). ParTI analyzes a continuum of gene expression according to specialist and generalist terms, where specialist cells are close to the vertices of a polytope (triangle, tetrahedron, and so on) that encloses the gene expression continuum. The main tasks of the population are inferred from the gene expression of cells closest to each of the vertices.

**Figure 3.**
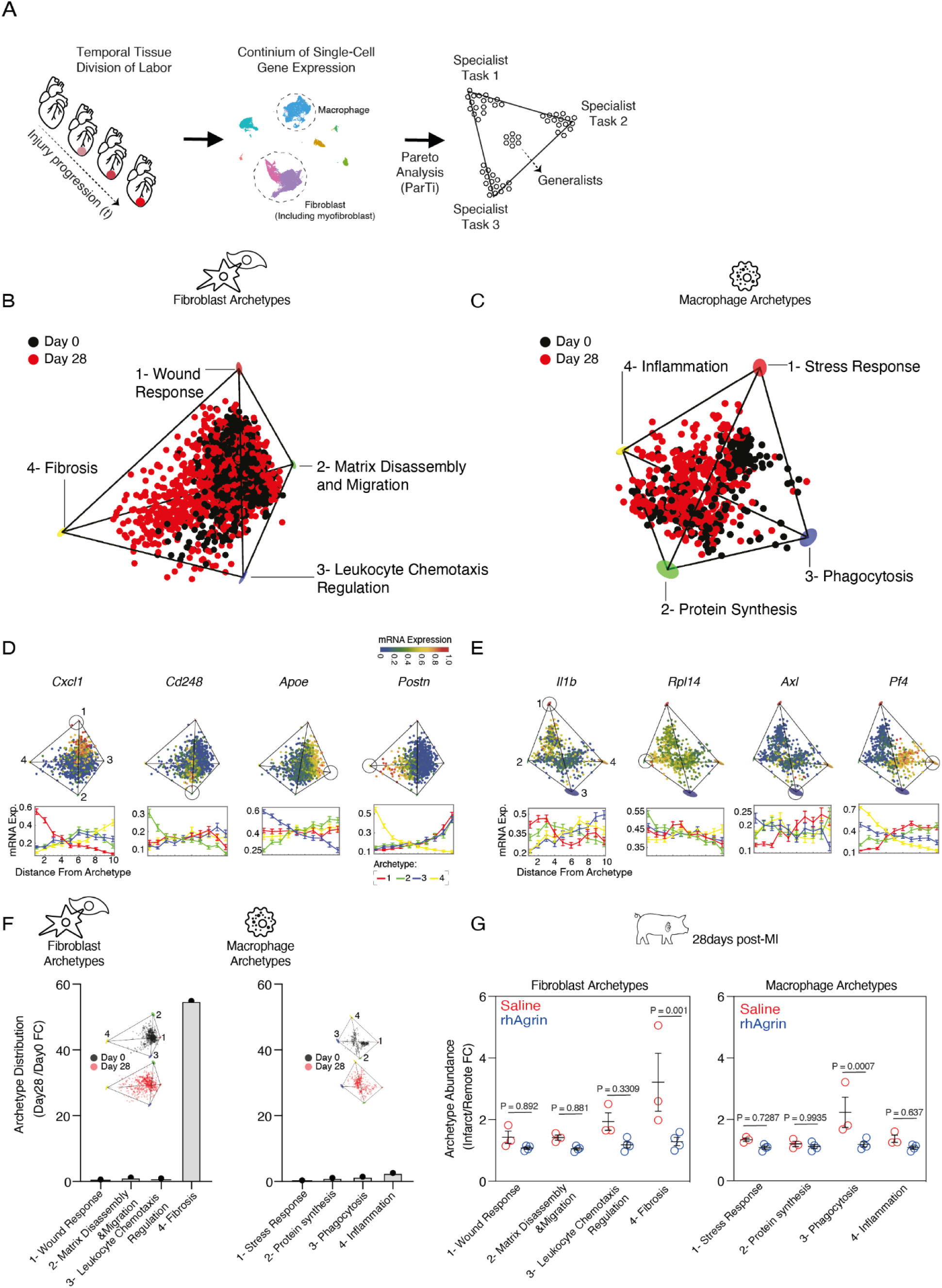
In cold fibrosis, macrophages return to homeostatic functions whereas fibroblasts acquire a fibrotic cell state. **(A)** schematic description of macrophage and fibroblasts temporal division of labor assessment after MI. Pareto analysis was used to characterize the continuum of gene expression within each cell type as distributed between specialist and generalist cells. Pareto analysis was done using ParTi (Hart et al., 2015). Fibroblasts (including myofibroblasts) **(B)** and **(C)** macrophages from day 0 (black) and day 28 (red) following MI fill in a tetrahedron in gene expression space (*P-value< 0.0001* for fibroblasts, *P-value= 0.02* for macrophages). The cells are projected on the first three principal components. **(D)** Gene expression of the top enriched genes for the fibroblast and **(E)** macrophage archetypes. Upper panel: cells within the tetrahedron are colored by the normalized mRNA expression level of each gene (on a scale of 0-1). Lower panel: mRNA expression level plotted as a function of the Euclidean distance from each archetype (in gene expression space) where cells were binned into 10 bins and the mean expression across the bins was computed. Archetypes for macrophages and fibroblasts are denoted based on color (1- red, 2- green, 3- blue and 4- yellow). **(F)** Fold change (FC) of proportion of cells from day 28 (red) and day 0 (black) that are closest to each archetype. **(G)** Deconvolution of Bulk mRNA sequencing of pig hearts following MI of either rhAgrin (blue) or saline treated (red) samples. Macrophage and fibroblast archetype abundances were assessed by gene signatures as fold change(FC), between infarct and remote zones (Methods, Table S7). Deconvolution of Bulk mRNA sequencing of pig hearts following MI of either rhAgrin (blue) or saline treated (red) samples. Macrophage and myofibroblast abundances were assessed by gene signatures as fold change(FC),between infarct and remote zones (Methods, Table S6). Statistical analysis was performed by 2-way ANOVA with multiple comparisons.

Macrophages and fibroblasts each showed 4 major tasks (*P-value<0.0001* fibroblasts, *P-value=0.02* macrophages) (**Fig. 3B-C**). The macrophage tasks relate to stress response, protein synthesis, phagocytosis, and inflammation, while the fibroblast tasks include wound response, matrix disassembly/migration, leukocyte regulation and fibrosis/ECM deposition, as determined by GO analysis (**Fig. 3B-E, fig. S3A-D, Table S7**). Interestingly, fibroblasts at day 28 post-MI almost exclusively populate archetype #4: the fibrosis/ECM deposition task (**Fig. 3F**). This indicates that during MI-driven cold fibrosis, fibroblasts acquire distinct cell state changes, consistent with a previous study (Fu et al., 2018). In contrast, macrophages are spread more equally between the 4 archetypes at both day 0 and 28 (**Fig. 3F**). This is consistent with our flow cytometry analysis suggesting that macrophage identity, activity and numbers return to baseline levels between days 14 and 65, when cold fibrosis is highly prominent (**Fig. 1D-G**). Interestingly, we find the gene signature of fibroblast archetype #4 to be highly enriched in pig saline treated infarct zones 28 days following MI (**Fig. 3G**). We conclude that in cold fibrosis macrophages return to a quiescent homeostatic state whereas fibroblasts persist and acquire a distinct pro-fibrotic phenotype, which is consistent across these two species.

### The circuit model predicts that the myofibroblast autocrine loop is a target for reducing cold fibrosis

We next asked which molecular interactions might be suitable targets for reducing cold fibrosis using *in-silico* perturbations of our mathematical model. We simulated different interventions by changing each of the model parameters. For each such changed parameter, we computed the basin of attraction to the healing state (**Fig. 4A**). The parameter with the largest impact is the myofibroblast autocrine growth-factor loop (Adler et al., 2020). According to the model, inhibition of the myofibroblasts autocrine loop significantly expands the healing margins. Notably, at a threshold level of about 40% inhibition of myofibroblast autocrine growth factor production, the cold fibrosis fixed-point vanishes, and the only steady state possible is the healing state where both myofibroblasts and activated macrophages return to baseline (**Fig. 4B**). Thus, our model indicates that the myofibroblast autocrine growth factor loop is a key target for reducing cold fibrosis. It further suggests that the inhibition need not be very strong, and even moderate inhibition of this loop can be beneficial.

**Figure 4:**
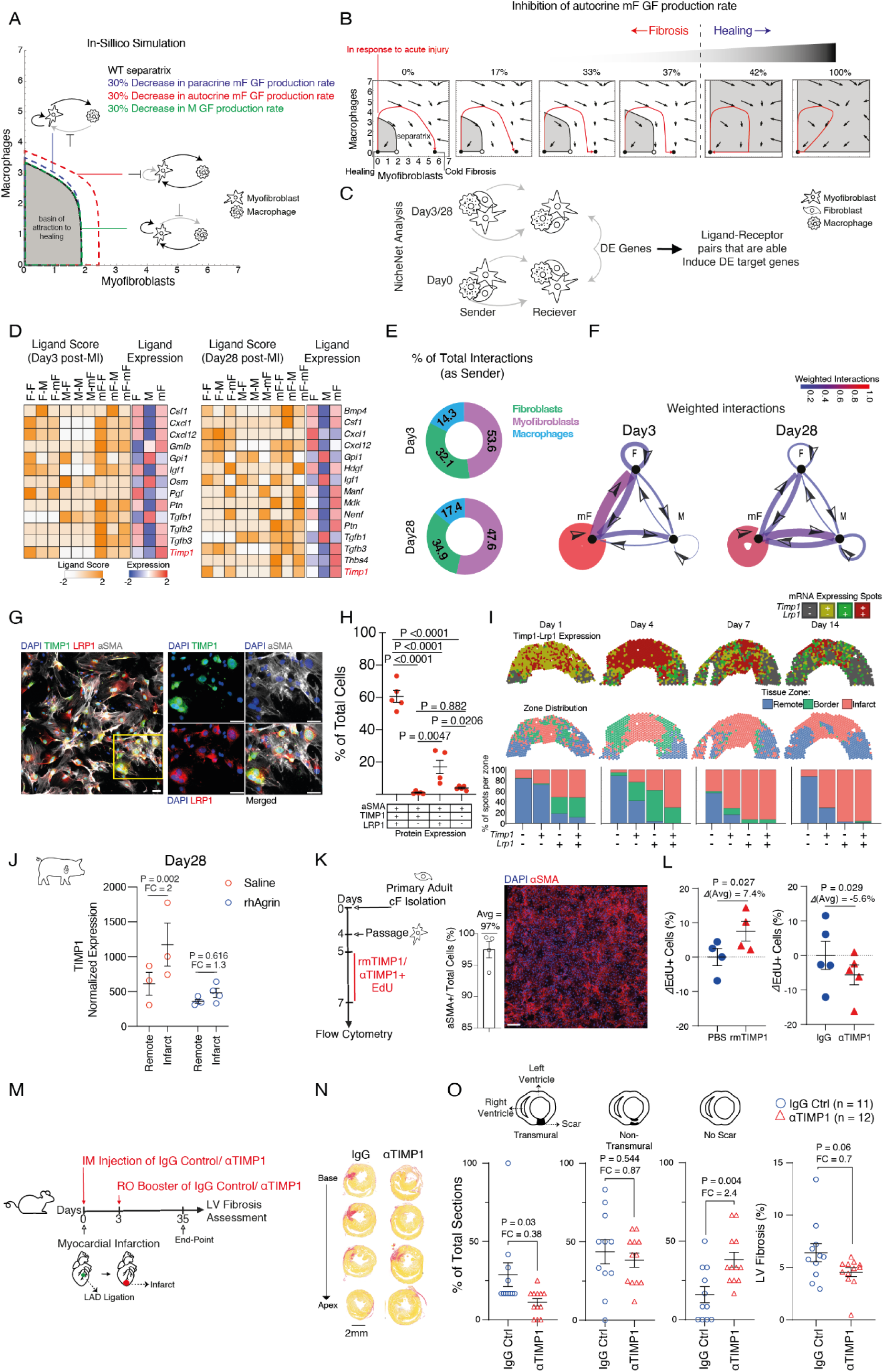
Targeting the autocrine loop of myofibroblasts by inhibiting TIMP1 reduces fibrosis after acute-MI in adult mice. **(A)** Phase plot where the axes represent the amount of cells. The separatrix (dashed lines) separates the basins of attraction of healing (gray area) and cold fibrosis (white). The separatrix is calculated with reference (WT) parameters (black), and with a 30% decrease in the production rates of the macrophage paracrine growth factor (green), myofibroblast paracrine growth factor (blue), and myofibroblast autocrine growth factor (red). **(B)** Simulations of the cell circuit response (red lines) to acute injury that leads to cold fibrosis with WT parameters (left panel). Healing is seen with a 42% or larger decrease in the myofibroblast autocrine growth factor production rate. **(C)** Schematic representation of NicheNet analysis to identify ligand-receptor interactions between fibroblasts (F), macrophages (M) and myofibroblasts (mF) at day 3 and 28 following MI. NicheNet performed on the scRNAseq data of Fig. 1B and fig. S1A-B. **(D)** Heatmap of potential ligands based on F, M and mF differentially expressed (DE) genes at days 3 (right) and 28 (left) post-MI. Ligand scores (Pearson correlation) and average ligand expression per cell type are presented as mean± SD (by either white-orange scale or blue-red scale, respectively). **(E)** Pie charts presenting the % of total interactions as sender per cell type and time point post-MI (day 3 or 28). Presented ligands are predicted by NicheNet analysis. **(F)** Weighted interactions (where interaction strength is normalized to the total number of sender and receiver cells) between F, M and mF following-MI (day 3 and 28) **(G)** Representative immunofluorescence images of TIMP1 (green), LRP1 (red), alpha smooth muscle actin (ɑSMA, grey) and DAPI (blue) in primary adult cardiac myofibroblasts cultures. (n = 5 biological replicates, 2510±515 SEM cells per replicate). Scale bars = 50μm, white frame represent the inset on the right. Quantification of images **(H)** based on their TIMP1, LRP1 and ɑSMA protein expression (as % of total cells) presented as mean± SEM. **(I)** Top panels- *Timp1* and *Lrp1* spatio-temporal distribution after MI. Positive spots (either *Timp1*, *Lrp1* or both) were defined as spots with expression, of each gene, higher than the median expression over all slices. Middle panels- zone distribution. Bottom panels- quantification of *Timp1*-*Lrp1* spot distribution, divided per zone. % of spots (based on *Timp1*-*Lrp1* expression) per zone are presented in a percentile graph. **(J)** Pig TIMP1 normalized mRNA expression at day 28 post-MI, in infarct and remote zones of rhAgrin (blue) and saline (red) treated hearts. Circles denote biological replicates. **(K)** Schematic of flow cytometry experiment for EdU incorporation and representative images of primary cardiac myofibroblasts cultures (day 4 of culture). Fraction of myofibroblasts at day 4 (n=4 relicates, mean cells=4253±749 SEM). Scale bar = 50μm. **(L)** Cardiac myofibroblast proliferation was assessed in cultures by means of delta EdU^+^ (%) cells of control (either PBS or IgG) and 48hrs treated cells (either rmTIMP1 or ɑTIMP1, respectively). In rmTIMP1 and ɑTIMP1 experiments n = 4 and 5 biological replicates, respectively). Data is represented as mean± SEM. Avg = Average. **(M)** Schematic experimental plan by which adult mice underwent MI and treated with either TIMP1 neutralizing antibodies (ɑTIMP1, n = 12 biological replicates) or IgG control (n = 11 biological replicates) immediately after injury onset and 3 days after injury. At 35 days left ventricular (LV) fibrosis was assessed by picro-sirius red staining. **(N)** Representative sequential picro-sirius red staining images along the the base to apex axis of ɑTIMP1/IgG control treated hearts. Scale bars: 2 mm. **(O)** Fibrosis parameters presented as % of total sections that are either transmural, non-transmural or sections with no scar (methods), and scar area out of the LV quantification.

### Timp1 is a cardiac myofibroblast autocrine growth factor

We therefore sought to characterize the growth factors (GF) that affect the myofibroblast autocrine loop. To do so we used NicheNet, an algorithm that infers ligand-receptor interactions within a transcriptomic dataset, using prior knowledge on ligand-receptor interactions and regulatory networks (Browaeys, Saelens, & Saeys, 2020). We applied NicheNet to the scRNAseq dataset by Forte, et al. (**Fig. 4C-D, fig. S4A**) (Forte et al., 2020). To focus on growth factor interactions, we only retained ligands according to a curated list of GFs (**Methods**). This analysis detects paracrine and autocrine growth factor interactions of macrophages (M), fibroblasts (F) and myofibroblasts (mF) (**Fig. 4D**). Among the growth factor interactions between the cell populations, the strongest is the autocrine loop of myofibroblasts in terms of interaction numbers and expression (**Fig. 4E-F**).

To establish whether each interaction is specific to the cold fibrosis state, we asked whether the interaction is differentially highlighted between day 0 and day 3 or 28 post-MI (**Fig. 4D**). This includes known factors previously shown to play important roles in cardiac fibrosis, such as Transforming Growth Factor beta (*Tgfb1, Tgfb2, Tgfb3*) (Stempien-Otero, Kim, & Davis, 2016; Tallquist & Molkentin, 2017) and Oncostatin-M (*Osm*) (Abe et al., 2019). We also find Tissue Inhibitor of Metalloproteinase 1 (*Timp1*) and Thrombospondin 4 (*Thbs4*) which were not previously characterized as cardiac myofibroblast growth factors.

We focused on *Timp1* because it was detected on both day 3 and 28 of the NicheNet analysis, and it demonstrated a myofibroblast autocrine signaling pattern. Timp1 has been studied in relation to fibrosis mostly in the context of its inhibition of matrix enzymatic activity as a pro-fibrotic and an anti-fibrotic factor (Grünwald, Schoeps, & Krüger, 2019; H. Wang et al., 2011). Following MI, *Timp1* is expressed almost exclusively by cardiac myofibroblasts (**fig. S4B**). Its expression peaks 3 days post-MI, coinciding with the peak of myofibroblast proliferation (**fig. S4C, H**). *Timp1* expression persists for 28 days after MI in line with the cold fibrosis phenotype (**fig. S4C**).

The predicted receptor for Timp1, the low-density lipoprotein receptor related protein 1 (Lrp1) is expressed by myofibroblasts and other cell types (**fig. S4A, E**). In myofibroblasts, *Lrp1* mRNA peaks at day 3 and persists at day 28 in scRNAseq analysis (**fig. S4F**).

The P1 (postnatal day 1) mouse heart is well known for its robust regenerative potential (Porrello et al., 2013). Bulk-mRNA analysis of hearts after MI shows a transient peak of *Timp1* and *Lrp1* expression in P1 hearts, while hearts of non-regenerative, P8, mice had sustained expression up to 7 days post-MI (**fig. S4D, G**). In pigs, TIMP1 mRNA expression is upregulated at day 3 and maintained 28 days after MI, but diminished following rhAgrin treatment (**Fig. 4J, Table S2-5**). To further test Timp1 and Lrp1 as an autocrine ligand-receptor pair, we co-stained them in primary cardiac myofibroblasts obtained from adult mouse hearts (**Fig. 4G-H**). The majority (60%) of cardiac myofibroblasts co-expressed TIMP1 and LRP1. A minority (20%) expressed LRP1 alone, with very few cells expressing TIMP1 alone or neither (**Fig. 4H**). Notably, our spatial transcriptomics Visium dataset shows that *Timp1*-*Lrp1* co-expressing spots become more abundant in the infarct zone over time (**Fig. 4I**).

To test the growth factor activity of TIMP1, we used an *in-vitro* culture assay of primary cardiac myofibroblasts (**Fig. 4K**) which were treated with either recombinant mouse TIMP1 (rmTIMP1) or with a neutralizing antibody against TIMP1 (ɑTIMP1). rmTIMP1 induced an increase in EdU (5-ethynyl-2’-deoxyuridine) incorporation in myofibroblasts, a marker of cell division, whereas inhibition of TIMP1 by the neutralizing antibodies reduced EdU incorporation (**Fig. 4L**). Based on this *in-vitro* culture assay we conclude that TIMP1 acts as a growth factor for cardiac myofibroblasts. It is expressed by cardiac myofibroblasts after injury together with its predicted receptor Lrp1, forming an autocrine signaling loop.

### Inhibiting Timp1 reduces fibrosis after acute cardiac injury

The mathematical model predicts that inhibition of the autocrine growth factor loop should reduce fibrosis (**Fig. 4A**). To test this *in-vivo*, we performed MI using permanent LAD ligation on adult mice. We then injected control (IgG) or ɑTIMP1 neutralizing antibodies directly to the injured myocardium following injury, and again 3 days after injury by retro-orbital sinus injection (RO) (**Fig. 4M**). We analyzed heart sections 35 days after injury (**Fig. 4N-O**). Mice treated with ɑTIMP1 antibody showed reduced fibrosis - the number of slides with detectable scarring was 2.4-fold lower in the ɑTIMP1 treated mice compared to mice treated with the control antibody. ɑTIMP1 treatment reduced the severity of the scar by 2.5-fold as measured by the number of sections that showed a transmural scar, defined as a scar that is continuous through the entire left ventricular wall. We conclude that inhibiting TIMP1 reduces fibrosis after MI.

## DISCUSSION

To investigate the concepts of hot and cold fibrosis, and to identify targets that could reduce fibrosis following acute cardiac injury, we combined mathematical modeling and *in-vivo* experiments in mice and pigs. According to the model, there are two types of fibrosis, one with both myofibroblasts and macrophages - hot fibrosis, and another driven by myofibroblasts alone- cold fibrosis. We show that the trajectory of acute MI goes towards cold fibrosis, in both mice and pigs. Fibroblasts acquire a profibrotic phenotype in cold fibrosis, and support their own growth by means of a growth factor autocrine loop. Using a circuit-to-target approach we predicted that inhibiting this autocrine loop should abrogate fibrosis. We identify TIMP1 as a cardiac myofibroblast autocrine growth factor. Pharmacological inhibition of TIMP1 with an antibody reduced fibrosis in adult mice following MI.

The present study highlights the utility of minimal mathematical models for identifying a target to reduce fibrosis. Targeting the autocrine loop makes sense because in cold fibrosis, the absence of macrophages leaves myofibroblasts as the primary source of their own growth factors. Importantly, the model predicts that in order to abolish the cold fibrosis fixed point, it is not necessary to inhibit the autocrine loop completely. Instead, the loop needs only to be inhibited below a threshold (∼40%) leaving a single regenerative healing fixed point.

In pigs we show that cold fibrosis occurs after acute MI, and that rhAgrin treatment reduces fibrosis and shifts gene expression towards that of an uninjured tissue. The concept of cold fibrosis after MI, is also supported by a recent analysis of human hearts samples showing high levels of activated fibroblasts and low inflammatory scores (Amrute et al., 2022).

We identified TIMP1 as a cardiac myofibroblast autocrine growth factor and as a target for reducing fibrosis after MI. TIMP1 is an inhibitor of metalloproteases (MMPs), although its activity as an MMPs inhibitor is lower than that of other TIMPs such as TIMP2, 3 and 4 (Brew & Nagase, 2010). Although TIMP1 is mainly discussed as an MMP inhibitor, it has also been described as a growth factor of several cell types, including scleroderma fibroblasts (Kikuchi, Kadono, Furue, & Tamaki, 1997) and hepatic stellate cells (Fowell et al., 2011). Importantly, TIMP1-associated cellular proliferation was observed independent of its MMP inhibition activity (Bertaux, Hornebeck, Eisen, & Dubertret, 1991; Hayakawa, Yamashita, Tanzawa, Uchijima, & Iwata, 1992). TIMP1 knockout mice showed reduced fibrosis in a model of hypertension-induced chronic cardiac injury (Takawale et al., 2018). In contrast, these TIMP1 knockout mice had an exacerbated acute liver fibrosis (H. Wang et al., 2011). Anti-TIMP1 antibody treatment reduced virus-induced myocarditis in mice (Crocker et al., 2007). It would be intriguing to see whether inhibiting TIMP1 might alleviate fibrosis in other contexts in which it acts as a growth factor for pro-fibrotic cells.

Our results indicate that the pathology of acute MI consists of two phases, which could be viewed as two distinct disease states. The early phase is characterized by the activation of both myofibroblasts and macrophages, which are also recruited from other tissues via the blood system along with other immune cells. Using spatial transcriptomics (Visium) we identified a distinct border zone population that separates the infarct from the healthy myocardium. Interestingly, the border zone seems to be reduced at the shift between the inflammatory phase to the cold fibrotic state (7-14 days following MI). The existence of these two phases is important to understand the pathophysiology of fibrosis following MI. It would be interesting to determine whether similar spatio-temporal tissue dynamics exist in other organs after injury. These dynamics are consistent with the notion that myofibroblast activation and the inflammation that occur after MI are part of the endogenous healing process; it is only when both processes are over-activated that detrimental consequences occur (Tzahor & Dimmeler, 2022).

The finding of cold fibrosis after acute MI raises the question of which other pathologies might show hot fibrosis. Recent histological analysis of human transplanted kidneys indicated hot and cold fibrosis states that are spatially-dependent (Setten et al., 2022). The same study extended the mathematical model (Adler et al., 2020) to include the effects of local hypoxia and inflammation status on the cell circuit. We hypothesize that scenarios of recurring injury and chronic inflammation might be especially prone to hot fibrosis. The cancer microenvironment might also share similarities to hot fibrosis, with extensive crosstalk between cancer associated fibroblasts and tumor associated macrophages (Adler et al., 2020; G. Friedman et al., 2020). Hot fibrosis may require different treatment strategies than cold fibrosis, due to the presence of activated macrophages that can support the growth of myofibroblasts (Adler et al., 2020).

Inhibiting the myofibroblast autocrine loop is an example of a more general strategy in which one can collapse a cell population by modulating growth factor interactions. This strategy differs in a qualitative way from other approaches that aim at affecting ECM deposition from pro-fibrotic cells or modulating their phenotypes. The collapse of a cell type removes all of its secreted factors in a single step, and thus may be more decisive as a treatment for a complex pathology such as fibrosis. An example is the recent use of Chimeric Antigen Receptor-T (CAR-T) technology to specifically attack cardiac myofibroblasts (Aghajanian et al., 2019). Killing of cells as occurs using the CAR-T technology can lead to excessive immune activation (Davila et al., 2014; Schmidts, Wehrli, & Maus, 2020).

In summary, we establish that cold fibrosis develops over time after myocardial infarction and use this concept to demonstrate a treatment strategy to reduce fibrotic scarring in the heart by inhibiting an autocrine growth factor of myofibroblasts. Our study highlights the utility of mechanistic yet simple mathematical models to cut through the complexity of pathological processes, to provide useful concepts to understand fibrosis and molecular strategies for treating this pathology.

## MATERIALS AND METHODS

### Mice

All mice used in this study were purchased from Envigo in the appropriate age [Hsd: ICR (CD1), Female, 31-34 grams, 10-12 weeks] for MI surgery. Mice arrived at least 4 days before the procedure and were housed in a pathogen-free environment at the animal facility, Weizmann Institute of Science. All experimental procedures were approved by the Animal Care Committee of the Weizmann Institute of Science (approval numbers: 00660121-2, 00510122-3).

### Pigs

Landrace pigs, 3 months old (male and female) ∼22-35 kg, were purchased from a local farm. Animal care and all experimental procedures were performed in strict accordance with the German and National Institutes of Health animal legislation guidelines and were approved by the local animal care and use committees.

### Mouse myocardial infarction

Myocardial infarction (MI) was induced by a permanent ligation of the left anterior descending (LAD) coronary artery. Adult ICR mice were anesthetized using isoflurane (Abbott Laboratories, Cat# AWN-34014202) and, following tracheal intubation, were artificially ventilated. Toe pinche was performed to test the depth of anesthesia. Prior to all surgical procedures mice were administered with buprenorphine in a subcutaneous manner, as an analgesic agent (0.066 mg/kg). Lateral thoracotomy was performed at the third intercostal space by blunt dissection of the intercostal muscles. Following the exposure of the left ventricle the LAD an 8-0 silk suture was used to tie the LAD coronary artery. The suture was validated by holding in the tie for 5 seconds, and the observance of an ischemic (white) area below the suture. Following this validation, the procedure ended by closing the chest cavity and skin by either suture and biological glue- Histoacryl® (B. Braun Melsungen AG, Cat# LUXBBS105005/2), respectively. Following awakening, mice were warmed for several minutes in a recovery cage.

For anti-TIMP1 neutralizing antibody experiment- immediately after LAD ligation, anti-TIMP1 (AF980, R&D) antibodies or IgG control (AB-108-C, R&D) were be injected using an insulin syringe directly to the myocardium of the infarct area at a concentration of 0.1ug/gr. Mice were also given a booster injection with a single retro-orbital sinus injection of anti-TIMP1 antibodies or IgG control (0.166ug/gr). For the evaluation of baseline cardiac function and after injury we used transthoracic electrocardiography performed on mice sedated with 2-3% isoflurane (Abbot Laboratories) using Vevo3100 (VisualSonics). Analysis was performed with Vevo Lab 3.2.6 software (VisualSonics). Mice with baseline ejection fraction (EF) values lower than 45% were excluded from the experiments. Mice were then randomized to treatment groups. Animals that were too severely injured 3 days following injury (EF< 30%) were excluded from the experiment.

### Pig myocardial infarction (ischemia-reperfusion injury)

All pig experiments were conducted in the Institute for Surgical Research at the Technical University of Munich (TUM) as described previously (Baehr et al., 2020; Hinkel et al., 2015). Briefly, a balloon was placed in the LAD distal to the bifurcation of the first diagonal branch and inflated with 6 atm. Correct localization of the coronary occlusion and patency of the first diagonal branch was ensured by injection of contrast agent. The percutaneous transluminal coronary angioplasty balloon was deflated after 60 minutes of ischemia; the onset of reperfusion was documented angiographically. At the onset of reperfusion, animals were treated with 33 µg/Kg (in 5mL of saline) of rhAgrin (#6624-AG, R&D Biosystems) or saline alone as control.

### Pig heart Samples

Pig heart samples were collected at day 3 and 28 following the MI. Hearts were harvested and stained by Tetrazolium chloride (TTC) to visualize scar/infarct. Left ventricles were dissected transversely to 5, 1cm thick slices, and each slice was further sectioned into 4/8 sections. Sections were annotated as infarct zones (positive for TTC) and remote zones (TTC negative). Sections were frozen on dry ice for further analysis, and were later subjected to RNA purification/ histological staining.

### Flow Cytometry

Adult mice were euthanized by neck dislocation and hearts were perfused with 10 ml of cold phosphate buffered saline (PBS). Left ventricles were dissected, chopped finely and digested in RPMI containing Collagenase-I (Sigma, 450U/ml), Collagenase-XI (Sigma, 120U/ml), Hyaluronidase (Sigma, 60U/ml) and DNase-I (Sigma, 10mg/ml), 37°C, for 1 hour, while agitated (200rpm). Digested materials were passed through a 100um metal mesh into 12 ml PBS and pelted by cooled centrifugation (4°C, 1400rpm, 5min). Supernatant was removed and the pellet was resuspended in 3ml of MACS buffer (2%FBS, 1mM EDTA in PBS). Cells were collected by cooled centrifugation (4°C, 1400rpm, 5min), supernatant was removed, and samples were blocked in 50ul of anti-CD16/32 antibodies (BioLegend, clone: 93, 1:200) in MACS buffer (20min, 4°C). Samples were resuspended in a 3ml MACS buffer, centrifuged (4°C, 1400rpm, 5min), supernatant was removed, and cell surfaces were stained in 50ul of antibody mix in MACS buffer (20 min, 4°C in the dark). Samples were washed in a 3ml MACS buffer containing DAPI (1:10,000) and pelted by cooled centrifugation (4°C, 1400rpm, 5min). Single cell suspension was resuspended in 0.5ml of MACS buffer and filtered through 40 um mesh prior to flow cytometry analysis by Fortessa (BD Biosciences, BD Diva Software).

For primary cardiac myofibroblasts proliferation assay, cells were incubation with 10uM EdU (5-ethynyl-2’ -deoxyuridine) (Invitrogen, C10424) for 48hours together with either recombinant-mouse TIMP1 protein (1µg/ml, ab206786, Abcam), PBS, anti-TIMP1 (1ug/ml, AF980, R&D) antibodies or IgG control (1µg/ml, AB-108-C, R&D). Cells were washed once with warm (37°C) PBS and then lifted from wells using Trypsin EDTA solution B (Sartorius, Cat# 03-052-1B) (5mins, 37°C). Trypsin solution B was neutralized by adding full-DMEM media and cells were pelted by centrifugation (Room temperature, 2200rpm, 5min). Cells were then washed once with 1% bovine serum albumin (BSA) in PBS and pelted by cooled centrifugation (4°C, 1400rpm, 5min). Cells were fixed and permed using Click-IT fixative and perm buffers and further processed based on the manufacturer’s instructions (Invitrogen, Cat# C10424). Cells were then filtered through a 40μm mesh prior to flow cytometry analysis. Following data acquisition single cell analysis was performed using the FlowJo (v10.6.2) software.

Gating for flow cytometry analysis: Myeloid cells were defined as Single cells, DAPI^-^CD45^+^CD11b^+^. Infiltrating cells (Neutrophils and Monocytes) were defined as single-cells, DAPI^-^CD45^+^CD11b^+^GR-1^+^ cells. Macrophages were defined as single-cells, DAPI^-^CD45^+^CD11b^+^GR-1^-^F4/80^+^ cells, and further parsed based on TIM4 in order to confirm tissue residency (Dick et al., 2019). Proliferating primary cardiac myofibroblasts were defined as Edu^+^ cells/ total single cells identified.

Mouse Antibodies: CD45 (BioLegend, clone: 30-F11, 1:100), CD11b (BioLegend clone: M1/70, 1:100), GR-1 (BioLegend, clone: RB6-8C5, 1:100), F4/80 (Bio-Rad, clone: Cl:A3-1, 1:80), TIM4 (BioLegend, clone: RMT4-54, 1:100).

### Primary cardiac myofibroblast cultures

For primary cardiac myofibroblast enriched cultures cells were first isolated from adult mice hearts using a neonatal dissociation kit (Miltenyi Biotec,130-098-373) and gentleMACS homogenizer (Miltenyi Biotec). Following euthanasia hearts were extracted, rinsed in PBS and chopped finely on a sterile surface using fine scissors. Chopped pieces were then transferred to GentleMACS C-Tubes (Miltenyi Biotec, Cat# 130-093-237) together with 2.5ml of the neonatal dissociation kit (Miltenyi Biotec,130-098-373) digestion buffer (prepared based on manufacturer’s instructions). GentleMACS C-Tubes were then incubated for 25mins at 37°C, and then spun using the GentleMACS homogenizer (Miltenyi Biotec) for 30secs (spin program: *’m_neoheart_01_01’*). GentleMACS C-Tubes were then incubated at 37 °C for 25mins (this step was repeated twice to achieve a total of 4 incubations and spins). Following the 4th incubation in 37 °C for 25mins, cells were spun twice. Cell suspension was then neutralized with pre-heated (37 °C) full-DMEM media- DMEM/F12 (Sigma, Cat# D6421) supplemented with L-glutamine (1%; Biological Industries, Cat# 03-020-1B), pyruvate (1%; Biological Industries, Cat# 03-042-1B), non-essential amino acids (1%; Biological Industries, Cat# 01-340-1B), penicillin, streptomycin, amphotericin B (1%; Biological Industries, Cat# 03-033-1B), horse serum (5%; Biological Industries, Cat# 04-004-1A) and FBS (10%; Biological Industries, Cat# 04-007-1). Cells were then pelleted by centrifugation (5mins, room temperature, 2200rpm). Resuspended in 1ml full-DMEM media, cells were filtered through a 100μm m mesh and cultured in gelatin-coated (0.1%; Sigma, Cat# G1393) 6-well plates. The next day cells were washed three times and media was replaced with fresh media. In the 2rd day following isolation cells were washed twice and media was replaced with fresh media. On day four, cells were split and seeded in 12-well plates (10^5^ cells/well) for flow cytometry analysis or in 96-well plates (either 5*10^3^ or 2*10^4^ cells/well) for immunofluorescence staining. Cells were then cultured for 48 hours and either fixed, by 4% paraformaldehyde (Electron Microscopy Sciences, Cat# 15710), and stained for immunofluorescence analysis or further processed for flow cytometry analysis. Throughout the experimental period cells were placed in an incubator (37°C and 5% CO_2_).

### Immunofluorescence

Cells in 96-well plates were fixed by 4% paraformaldehyde for 10 minutes, followed by 3 washes of PBS (Room temperature) on a shaker. Following fixation cells were permeabilized by 0.1% Triton X-100 for 7 minutes and blocking with 2% BSA in PBS for 1 hour in room temperature, on a shaker. Primary antibodies were prepared in blocking solution and antibodies were incubated overnight (4°C, gentle agitation). Cells were then washed 3 times for 10 min with PBS and stained with the suitable secondary antibody (1:200, Jackson ImmunoResearch) and DAPI (4,6-diamidino-2-phenylindole dihydrochloride) for 1h at room temperature. Wells were later washed 3 times with PBS in room temperature, on a shaker. Images were obtained with a Nikon Eclipse Ti2 fluorescent microscope with Nikon NIS-Elements software.

The following primary antibodies were used for staining: anti-LRP1 (1:100, ab92544, Abcam), anti-TIMP1 (1:50, AF980, R&D systems), anti-ɑSMA (1:400, A2547, Sigma).

The following secondary antibodies were used for staining: Cy™3 AffiniPure Fab Fragment Goat Anti-Mouse IgG2a (Jackson ImmunoResearch, Cat# 115-167-186), Cy™3 AffiniPure Donkey Anti-Rabbit IgG (H+L) (Jackson ImmunoResearch, Cat# 711-165-152), Alexa Fluor® 647 AffiniPure Donkey Anti-Mouse IgG (H+L) (Jackson ImmunoResearch, Cat# 715-605-150), Donkey Anti-Goat IgG H&L (Alexa Fluor® 488) (ab150129).

### Cell quantification in culture

Following staining for LRP1, TIMP1, αSMA and DAPI, Images were obtained with a Nikon Eclipse Ti2 fluorescent microscope with Nikon’s NIS-Elements software and analyzed using QuPath v0.3.0 software. Nuclei detected based on DAPI staining. An object classifier was trained on representative images to identify positive signals for each of the markers that were used for staining.

### Histology and picro-sirius red staining

Mouse hearts were briefly perfused with 10ml cold PBS and then fixed with 4% paraformaldehyde (Electron Microscopy Sciences, Cat# 15710) (overnight, shaking, 4°C). Paraformaldehyde fixed hearts samples were cut in the middle in a transverse manner and both half-heart pieces representing Base and Apex were embedded together in paraffin (Leica, Cat# 39601006). Embedded hearts were cut into 6 rows (holding 12 sections, 5μm thick), separated by 300μm jumps in-between. Following block cutting, sections were baked overnight (37°C). Picro-sirius red staining was done based on manufacturer’s protocol (Abcam, Cat# ab246832). Picro-sirius red sections were imaged in bright-field mode using the PANNORAMIC SCAN II slide scanner (3DHISTECH). Images were processed by CaseViewer v2.4 (3DHISTECH).

### Fibrosis analysis

Scarred tissues were quantified in a blinded manner, based on picro-sirius red staining of serial cardiac sections spanning through the entire heart (base to apex). In-house, ImageJ script was used to assess the average percentage of fibrotic area out of the left ventricle. Mice transverse heart sections were also subjected to scar severity classification as previously shown (Shakked et al., 2022). Each section was classified to one out of three classes: transmural scar (scar that crossed the entire left ventricular wall), non-transmural scar (scars on a single or both sides of the ventricular wall that does not cross) or no scar. The number of sections for each category was represented as a percentage of total sections on the slide.

### Spatial transcriptome processing

Following MI, adult mice were euthanized, and hearts were harvested on days 1, 4, 7 and 14 post-Injury. Hearts were briefly washed in Optimal Cutting Temperature (OCT) (Sakura Finetek, Cat# 4583) solution and placed in a new mold containing OCT. The molds were placed in pre-cooled Isopentane bath in liquid N_2_. Molds were placed on dry ice until storage in -80°C. For tissue sectioning, blocks were placed in cryostat (Epredia CryoStar NX70 cryostat, Thermo Fisher) until equilibration and were sectioned at 10μm thickness. Blocks were cut and sections were placed on SuperFrost plus slides (Thermo Scientific, Cat# 630-0951) and were rapidly stained with H&E (quick wash with H_2_O, 15sec in Hematoxylin (Cat# 6765002, Fisher Scientific), quick wash with H_2_O, counterstain for 15sec with Eosin (Cat# 10483750, Fisher Scientific) and wash with H_2_O before mounting the slide) and evaluated for localization and extent of damage. This process was repeated until the optimal region was reached.

To assess optimal permeabilization conditions for Visium experiment, 8 adjacent sections were placed on the Visium Spatial Tissue Optimization Slide (Cat# 1000193, 10X Genomics) and were subjected to different permeabilization times. Optimal permeabilization time was 30mins and was used in the Visium experiment.

Once all sections were selected for Visium, blocks were placed in cryostat until equilibration along with the Visium spatial gene expression slide (10X Genomics, Cat# 1000187). Blocks were then aligned and a single 10μm section was placed on the Visium slide. After all sections were placed on the Visium slide, the downstream pipeline was continued in accordance with the manufacturer’s guidelines (10x genomics, Visium Spatial Gene Expression Reagent Kits User Guide). The slides were imaged with Axio Imager.Z2 (upright) with Zeiss Axiocam 506 Color camera. Visium data was processed with SpaceRanger (v1.3.0) with the images manually aligned, with reference genome mm10. The filtered feature barcode matrix of each slide was imported to Seurat (v4.1.1) and merged.

### Spatial transcriptomics analysis

To filter out poor quality spots, as well as spots localized in blood vessels and not overlapping cells, fractions of hemoglobin genes out of the total UMI reads was calculated for each spot. Spots with a hemoglobin fraction above 1% were removed. Next, using Loupe Browser (6.1.0), spots localized at the outer layers of the tissue (inside and out), were manually annotated as spots distanced up to 2 spots from tissue edges, and removed from analysis. Weakly expressed genes, with a total of less than 1000 UMI over all slides were ignored. UMI counts of the remaining spots were normalized by dividing each count to the spot’s UMI total, resulting in fractions of UMIs. This dataset of ∼7000 spots over all slices and ∼4500 genes was used for differential gene expression. Counts were further normalized using log normalization, and non-linear dimension reduction was carried out using the UMAP algorithm using default parameters. Unbiased clustering was performed using the Mathematica function *’FindClusters’* with algorithm ’MeanShift’ and sub option *‘NeighborhoodRadius’* set to 0.45. In order to further visualize the clusters on the first 2 components of the projection, smooth kernel distribution was calculated (‘SmoothDensityHistogram’) and density contours were plotted. The clustering resulted in 3 clusters which were projected back on the slices from the 4 time points, and identified as the three distinct regions of remote, border and infarct zones. Gene signatures for the border and infarct zones were selected by calculating fold change (FC) expression of all genes and selecting genes with high FC relative to the expression in the remote zone (internal control per slice/time point). Mann-Whitney test with Bonferroni correction was used to calculate statistical significance. For the sake of visualization each slice was rotated so that the infarct zone median (defined by infarct zone spots) is positioned at the top, and log efxpression was rescaled (between 0 and 1) over all slices so comparison between time points is valid. Double positive spots (e.g., *Postn^+/^Cd68^+^* in Fig.1L-M and *Timp1^+/^Lrp1*^+^ Fig. 4I) were identified as spots with expression higher than the median expression over all slices.

### Pig mRNA library preparation and sequencing

Total RNA was extracted from 28 frozen pig samples in a paired manner (Remote and Infarct zones from the same animal were used to obtain optimal internal control [Agrin day 3 (n = 4), Saline day 3 (n = 3), Agrin day 28 (n = 4), Saline day 28 (n = 3)]. Following acquisition, frozen samples were crushed into fine powder using a mortar and pestle and further processed using miRNeasy Mini Kit (Qiagen, Cat# 217004) according to manufacturer’s guidelines. Replicates of high RNA integrity (RIN), determined by TapeStation (Agilent) (RIN=8.2±0.1) were processed for RNA-seq library preparation.

RNA-seq libraries were prepared at the Crown Genomics institute of the Nancy and Stephen Grand Israel National Center for Personalized Medicine, Weizmann Institute of Science. Libraries were prepared using the INCPM-mRNA-seq protocol. Briefly, the polyA fraction (mRNA) was purified from 500ng of total input RNA followed by fragmentation and the generation of double-stranded cDNA. After Agencourt Ampure XP beads cleanup (Beckman Coulter), end repair, A base addition, adapter ligation, and PCR amplification steps were performed. Libraries were quantified by Qubit (Thermo fisher scientific) and TapeStation (Agilent). Sequencing libraries were constructed with barcodes to allow multiplexing. Sequencing was done on a NovaSeq600 instrument (Illumina) using an S1 100 cycles kit, allocating approximately ±35.7million single-end 100-base-pair reads per sample (single read sequencing). Fastq files for each sample were generated by the usage bcl2fastq v2.20.0.422.

### Pig Bulk-mRNA sequencing analysis

Poly-A/T stretches and Illumina adapters were trimmed from the reads using cutadapt (Martin, 2011); resulting reads shorter than 30bp were discarded. Reads for each sample were aligned independently to the *Sus-Scrofa* reference genome Sscrofa11 using STAR (2.7.3a) (Dobin et al., 2013), supplied with gene annotations downloaded from Ensembl (release 103). The EndToEnd option was used and outFilterMismatchNoverLmax was set to 0.04. The percentage of the reads that were aligned uniquely to the genome was 90%. Deduplication of reads with the same UMI was performed by the PICARD (2.22.4) MarkDuplicate tool (using the BARCODE_TAG parameter). Expression levels for each gene were quantified using htseq-count (version 0.11.2) (Anders, Pyl, & Huber, 2015), using the gtf above. Only uniquely mapped reads were used to determine the number of reads falling into each gene (intersection-strict mode). Differential analysis was performed using DESeq2 package (1.26.0) (Love, Huber, & Anders, 2014) with the betaPrior, cooksCutoff and independentFiltering parameters set to False. Raw P values were adjusted for multiple testing using the procedure of Benjamini and Hochberg. Pipeline was run using snakemake (Köster & Rahmann, 2012). Differentially expressed genes were determined by a p-adj of < 0.05 and absolute fold changes > 2 and max raw counts > 30.

Hierarchical clustering (distance: Pearson’s dissimilarity, method: Ward.d) was performed, based on the 1000 most variable genes per condition [day (3 or 28)+treatment (agrin or saline)]. Clustering analysis was performed with Rstudio v3.6.1.

Unsupervised analysis was executed in order to explore the pattern of gene expression by clustering the genes based on 5961 genes (day 3) and 1079 genes (day 28). These genes were determined as differential expressed (DE) genes (upregulated or downregulated) in comparisons of infarct zones and remote zones in day 3 and 28 post-MI, in each of the treatment conditions (agrin and saline). Standardized, log2 normalized counts were used for the clustering analysis. Heat maps were constructed using Morpheus (https://software.broadinstitute.org/morpheus).

### Deconvolution of bulk-mRNA sequencing

To deconvolve our Pig bulk-mRNA data, raw counts were normalized per repeat to calculate the fraction of each gene from the total counts per sample. Repeats were averaged per gene per condition (Agrin or Saline), per time point (Day 3 and 28), and region (remote and infarct zones). Fractions of myofibroblasts/macrophages per time point, condition and zone were calculated by taking the average of the expression of marker gene lists for each cell type (macrophages, myofibroblasts) (**Table S6**). Fold change of cell fractions in the infarct zone per condition (Agrin, Saline) compared to the internal control (remote zone) were then calculated per time point (day 3 and 28). Finally, deconvolution of Pig bulk-mRNA data based on the mice archetypes was similarly done but with the signatures gene list extracted from the Pareto analysis (4 archetypical signature gene lists per cell type, Table S7). All calculations were done in Wolfram Mathematica 13.1. Signature gene list for myofibroblasts (88 genes), and macrophages (102 genes) were modified from the re-analyzed scRNAseq data of Forte, et al. (Table. S6) (Forte et al., 2020).

### Re-analysis of mouse Bulk-mRNA sequencing

Raw counts matrix of GSE123863, generated by (Z. Wang et al., 2019)) was downloaded from the GEO database. Genes with maximum counts of 5 or lower were filtered out and then differential expression analysis was done DESeq2 (Love et al., 2014) with the ‘*betaPrior’*, ‘*cooksCutoff’* and ‘*independentFiltering’* parameters set to False. Raw P values were adjusted for multiple testing using the procedure of Benjamini and Hochberg. Genes with average(logFC)>=1, p.adj<0.05 and above 30 counts in at least one of the samples, were considered as differentially expressed.

### Re-analysis of mouse single-cell mRNA sequencing

scRNAseq of cardiac interstitial cells from 7 different timepoints following MI (Forte et al., 2020) was downloaded from Array Express repository (E-MTAB-7895), and re-analyzed using Seurat V3.241. Cells were filtered according to 4 Median absolute deviation (MAD) below and above the median for each parameter (*‘nFeature_RNA’*, *‘nCount_RNA’*, *‘percent*.*mt’*) and sample. For % of mitochondrial reads, if the max % using the MAD is above 25% the last was used for filtering. We have also filtered out genes that appeared in less than 10 cells. Counts were normalized using log normalization. Next, non-linear dimension reduction was carried out using uniform manifold approximation and projection (UMAP) algorithm with the first 19 PCs of the data as input, using default parameters. Clusters were defined using *‘FindNeighbors’* and *‘FindClusters’* functions with 19 principle components (PCs) and resolution set to 0.5. Cluster annotation was done manually by visualizing the expression of known cell type specific marker genes and the top differentially expressed genes per cluster (*‘FindAllMarkers’* with the parameters: *logfc*.*threshold* = 0.5, *min*.*diff*.*pct* = 0.5). Calculations of macrophage and myofibroblast gene-signature enrichment was performed by *‘AddModuleScore’*. UMAP figures were produced by *‘FeaturePlot’* and dot plots of enriched genes was produced by Seurat’s *‘DotPlot’* function.

### Pareto analysis

We analyzed fibroblasts and macrophages scRNAseq data (Forte et al., 2020) and performed Pareto Optimality analysis to infer the trade-offs and tasks that the cells specialize in. The raw data includes 17,202 genes, 13,100 macrophages and 20,746 fibroblasts from 7 different time points following MI. We next consider cells from day 0 and day 28 only. Before preprocessing the data, we randomly sampled 500 fibroblasts from each time point such that we end up analyzing 1000 fibroblasts from days 0 and 28. For macrophages, we considered all 324 cells from day 0 and 401 cells from day 28 amounting to 724 cells. We then used ‘Sanity’ – a recently developed method to normalize single-cell data and to infer the transcription activity of the genes. Sanity is a unique Bayesian procedure for normalizing scRNAseq data from first principles (Breda, Zavolan, & Nimwegen, 2021). Following the Sanity normalization, we removed genes with low mean expression and low variation and considered genes with log10 mean expression larger than -14 and standard deviation larger than 0.03. This left us with 6007 genes for fibroblasts and 7028 genes for macrophages. We next used the ParTi package in Matlab (Hart et al., 2015) to fit the data of each cell type population to a polytope. We find that fibroblasts fill in a tetrahedron (P-value<0.0001) and macrophages are also best described by a tetrahedron with 4 archetypes (P-value=0.02). We next used ParTi to find enriched genes for all the archetypes in order to infer the tasks the cells specialize in. In order to infer specific pathway enrichment of either macrophage or fibroblasts tasks we used the enriched genes in each of the vertices. We specifically used pathway gene sets from biological processes of Gene Ontology (GO) (all GO analysis was done using the Mathematica Package MathIOmica (Mias et al., 2016).

To evaluate the distribution of the cells from day 0 versus day 28 within the tetrahedron, we sought to represent every cell as a convex combination of the archetype: 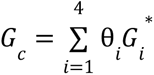 where *G*_c_ represents the gene expression of cell 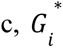 is the position of archetype i in gene expression space, and we are looking for θ*_i_* such that 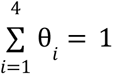. We then categorize the cells based on the archetype they are closest to (or its largest θ*_i_*). To compare the distribution of the cells from day 0 and day 28, we compared the proportion of the cells closest to each archetype from the total population per time point.

### NicheNet analysis

The re-analyzed scRNAseq by Forte et al., (2020) was used for Ligand-Receptor analysis by the NicheNet algorithm (Browaeys et al., 2020). The pairwise interaction between cell types (macrophages, fibroblasts, myofibroblasts) was calculated independently for each day (day 3 and 28 post-MI). For each cell type and for each day (day 3 and 28), genes of interest (GOIs) were defined genes that were upregulated between that day (3 or 28) and day zero and expressed in at least 25% of the cells in that cluster. Interaction score was calculated as the Pearson coefficient calculated by *‘predict_ligand_activities’* function, using the GOIs of that day and the genes that were expressed in the ‘receiver’ clusters as targets. Results were then refined by filtering against a list of growth factors (GO:0008083). Weighted interactions were calculated by considering the interaction score normalized to the number of interacting cells.

### *In-silico* myofibroblasts proliferation score

To estimate proliferation in myofibroblasts, we considered the expression of 98 mouse cell cycle genes (Tirosh et al., 2016). We considered the fraction of RNA counts of these cell cycle genes out of the total RNA counts in the myofibroblast cells considering cells that have total count between 10^3^ and 10^4^ molecules. To determine the difference between the distributions of proliferation scores from the different time points, we used a Mann-Whitney test to test whether the data is sampled from different distributions. Effect size (ES) was calculated based on the mean and standard deviation of each distribution where: 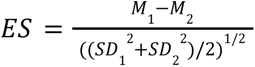.

### *In-silico* mathematical perturbations of cold fibrosis

We computed the effect of various potential targets on the circuit dynamics by comparing the outcome of weakening three growth factor interactions: the two paracrine interactions between myofibroblasts and macrophages and the autocrine growth factor interaction within the myofibroblast population. We consider the following mathematical model (based on the model presented in (Adler et al., 2020):

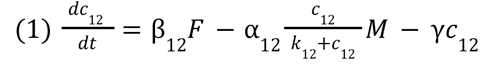

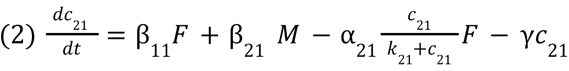

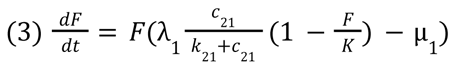

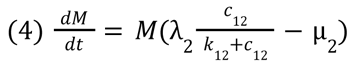

Where F, M are the concentration of fibroblasts and macrophages, and c_12_, c_21_ are the concentration of the two growth factors produced by the cells. We consider the following parameter values for the wild-type circuit:

Growth factor production rates:

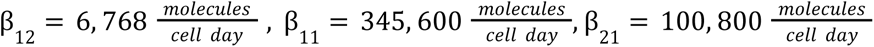

Growth factor endocytosis rates:

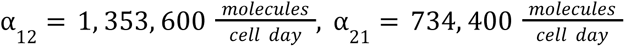

Growth factor degradation rate:

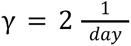

Growth factor binding affinities:

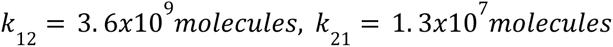

Cell maximal proliferation rates:

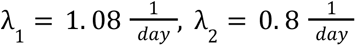

Cell death rates:

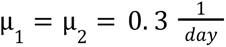

Carrying capacity of fibroblasts: *K* = 10^6^ cells

Most of the parameter values were determined based on Zhou, et al. (Zhou et al., 2018), with few exceptions. We consider the production rate of the macrophage growth factor, β_12_, to be 100-fold smaller in order to simulate a situation where the cold fibrosis state is a stable state (and there is no hot fibrosis). The macrophage growth factor binding affinity, *k*_12_, is considered to be 200-fold larger in order to correct for spatial effects.

To consider the effect of weakening growth factor production rates on the circuit dynamics, we compare the circuit with the wild-type parameters to three scenarios. In those three scenarios, we keep all model parameters at the same value and only decrease each of the production rates β_12_, β_21_, or β_11_ by 30%. To plot the separatrix, we numerically solve the final steady state levels of the cells (eqs. 1-4) where we sample different initial conditions.

To exemplify the effect of weakening the fibroblast autocrine loop (**Fig. 4B**), we considered the circuit (eqs. 1-4) with the wild-type parameters where we consider the decrease in the maximal proliferation rate of fibroblasts, λ_1_ .

### Statistical analysis

All experiments were carried out with n ≥ 3 biological replicates [except for Visium data produced from n = 1 per time point (day 1, 4, 7 and 14)]. Experimental groups were balanced for animal age, sex and weight. Different experimental groups were caged together and treated in the same way. Statistical analyses were carried out using the GraphPad Prism software (version 8.0.1). When comparing between two conditions, data were analyzed using a two-tailed Student’s t-test (Unpaired- Fig. 2B-C, Fig. 4O, Paired- Fig. 4L). When comparing more than two conditions, we employed an ANOVA analysis with multiple comparisons. Individual data points represent data derived from different biological repeats to demonstrate the distribution of data across different experiments. Statistical analysis of bulk-mRNA sequencing data is described under the appropriate methods subheadings. Statistical analysis is derived from the biological repeats of an experiment. Measurements are reported as the mean and the error bars denote the S.E.M. throughout the study (unless mentioned differently in figure legends). P-values are presented on figures in the appropriate comparison location. Statistical analysis of Pareto archetype inference and enrichment of genes was done using the ParTi package (Hart et al., 2015) in Matlab (R2022a). Statistics of myofibroblast proliferation score distributions were computed using the function *‘MannWhitneyTest’* in Mathematica 13.1.

### Illustrations

All figure Illustrations included in this study are original.

## Supporting information

Table_S4_DEoutput_D28_Agrin_IZvsRZ

Table_S2_DEoutput_D3_Agrin_IZvsRZ

Table_S5_DEoutput_D28_Saline_IZvsRZ

Table_S3_DEoutput_D3_Saline_IZvsRZ

Table_S1_Visium_DE

Table_S7_Macrophage_Fibroblast_Archetypes_Enriched_Genes

Table_S6_Macrophage_Myofibroblast_Gene_Signature

## Acknowledgements

We thank Marina Cohen from the Histology Unit, Department of Veterinary Resources, Weizmann Institute of Science, for histological processing and staining. We also thank Dr. Hadas Keren-Shaul and Danna Robbins (The Nancy & Stephen Grand Israel National Center for Personalized Medicine, Weizmann Institute of Science) for pig Bulk-mRNA sequencing library preparation and sequencing. We would like to acknowledge Histology and Next generation sequencing facilities at the Vienna Biocenter Core Facilities (VBCF) for all aspects of Visium handling, sectioning and sequencing. We thank Prof. Steffen Jung, Dr. Jung-Seok Kim and Sebastien Trzebanski from the Department of Immunology and Regenerative Biology, Weizmann Institute of Science, Rehovot, Israel, for insightful comments and technical assistance in flow cytometry experiments.

## Author contribution

Conceptualization: SM, MA, AM, UA, ET

Methodology: SM, MA, AM, EB

Investigation: SM, MA, AM, EB, YD, KBU, JE, AG, DK, DL, TS, MG, AS, LZ, JW, AB, CK

Visualization: SM, MA, AM, DMK

Funding acquisition: ET, UA

Project administration: ET, UA

Supervision: ET, UA, RM, EMT, CK

Writing – original draft: SM, MA, AM, UA, ET

Writing – review & editing: All authors

## Data and materials availability

All data are available in the main text or the supplementary materials. All count matrices and metadata for each transcriptomic dataset (Pig bulk-RNA sequencing and Visium spatial transcriptomics) will be publicly available in the Gene Expression Omnibus (http://www.ncbi.nlm.nih.gov/geo/) under data accession no. GSE###. All custom generated code will be available in the AlonLabWIS git.

---

**Figure S1:**
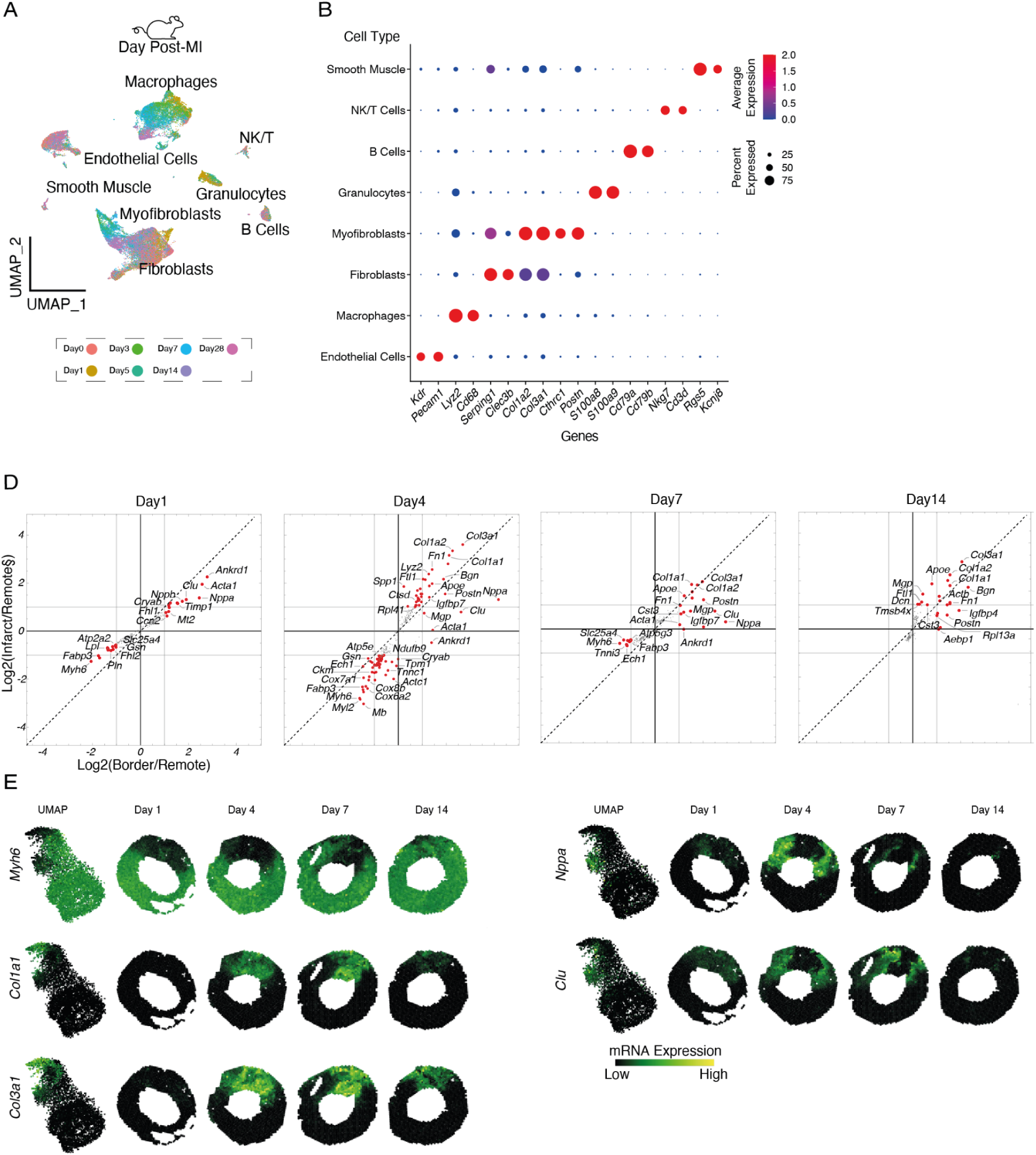
Single-cell mRNA-sequencing, and spatio-temporal differential gene expression analysis (Visium) following MI. **(A)** Complementary scRNAseq data in Fig. 1B. Total cardiac left ventricle interstitial cells following MI at days: 1 (yellow), 3 (light green), 5 (dark green), 7 (blue), 14 (purple) and 28 (pink). n = 1 per time-point. **(B)** Known and top differentially expressed genes per cluster used for cluster annotation. **(C)** Spatio-temporal differential gene expression analysis of adult mice following-MI. Log2 fold-change (FC) was calculated per area in comparison to the corresponding remote zone per section (used as internal control) (Infarct/Remote, Border/Remote). Differentially expressed genes (red spots) were defined as log2FC > 1 and significant p-values (<0.05) following Bonferroni correction. Small black dots represent non-differentially expressed genes. **(D)** Visium spatial cluster defining genes (Remote zone- *Myh6*, Infarct zone- *Col1a1*, *Col3a1*; Border zone- *Nppa*, *Clu*) projected on all sections and combined UMAP. mRNA expression is denoted by color (black to yellow). Each slice was rotated so that the infarct zone median (defined by infarct zone spots) is positioned upwards.

**Figure S2:**
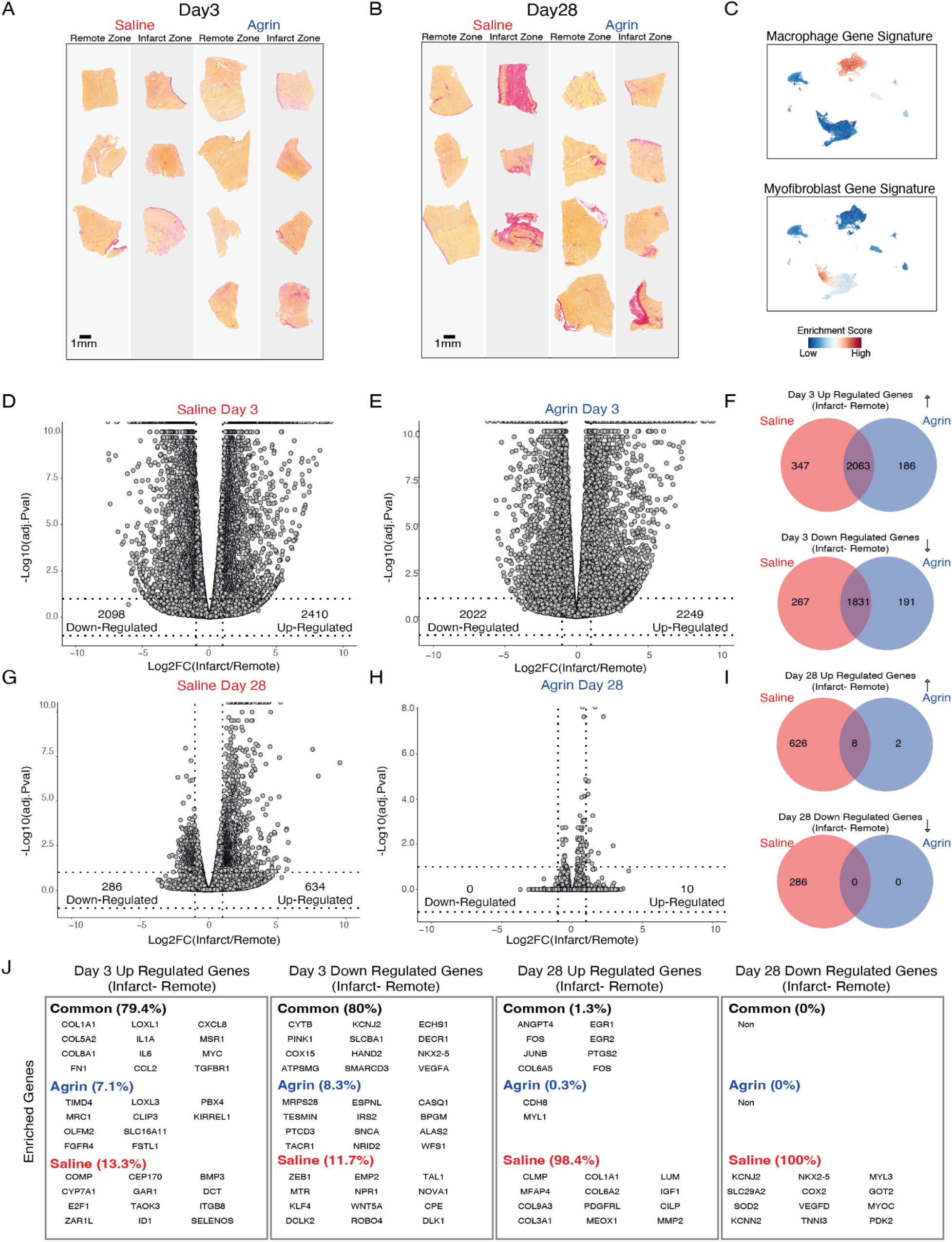
Differentially expressed genes in saline or rhAgrin treated porcine hearts following-MI. Representative picro-sirius red staining images of remote and infarct zones, from all saline and rhAgrin treated hearts from day 3 **(A)** and 28 **(B)** post-MI used for fibrosis quantification (Fig. 1B-C). Scale bars: 1 mm. **(C)** Validation of cardiac macrophage and myofibroblast gene signatures (Table S6) used for deconvolution analysis in Fig. 2G-H. Gene signature enrichment score was calculated for each gene set (cardiac macrophage and myofibroblast) and projected on a UMAP. Scores are presented from low to high and denoted by color (blue to red). Volcano plots of differentially expressed genes (annotated genes that correspond with the following conditions: |log2FoldChange| >= 1, p-adjusted value < 0.05 and max raw counts > 30) between remote and infarct zones of day 3 Saline **(D)**, rhAgrin **(E),** day 28 Saline **(G)** and rhAgrin **(H)**. Venn diagrams of common and unique up and down regulated genes between rhAgrin and saline treated hearts from day 3 **(F)** and 28 **(I)** post-MI. Numeric values represent the number of differentially expressed genes that are enriched exclusively in saline (red), agrin (blue) or both (overlapping red-blue region). **(J)** Highlighted genes for each Venn diagram analysis. % Are the amount of genes presented in each category (common, agrin or saline) calculated from all differentially expressed genes in the corresponding Venn diagram.

**Figure S3:**
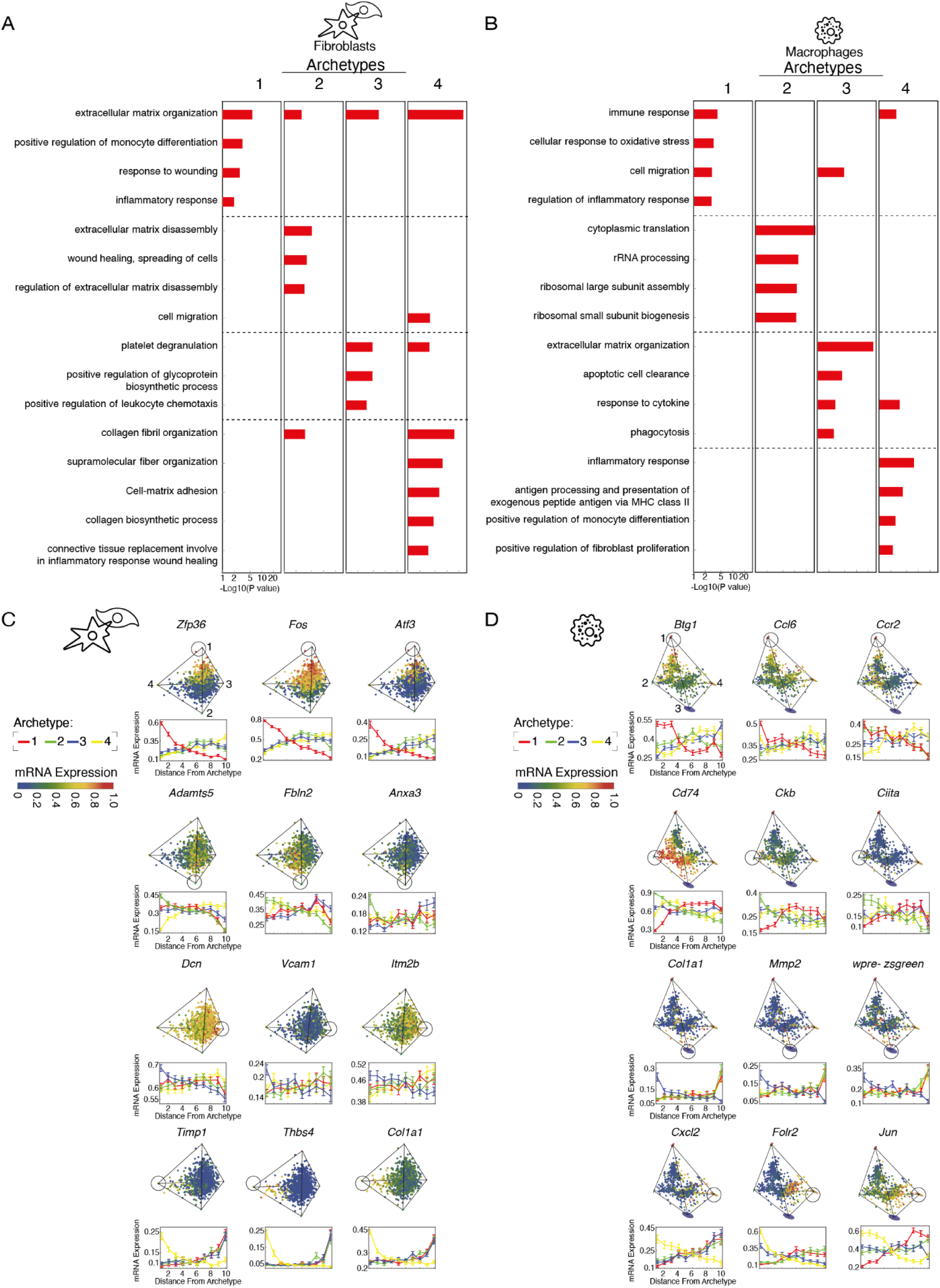
Archetype enrichment analysis. Table of top enriched GO terms for the different fibroblast **(A)** and macrophage **(B)** archetypes and their P-values in log scale. Examples of top enriched genes per fibroblast **(C)** and **(D)** macrophage archetypes. The expression of each gene is plotted in two ways: upper panels- the cells are colored based on the expression levels within the tetrahedron where they are projected on the first three principal components; lower panels- expression is plotted across 10 bins as a function of the Euclidean distance from each archetype in gene expression space. Red, green, blue, and yellow correspond to archetype numbers 1-4, respectively. **S3 supplementary text:** We note that archetype 3 in macrophages (**fig. S3D**), associated with phagocytotic functions, shows expression of prominent genes usually expressed in fibroblasts (*Col1a1*). It also shows expression of an epicardial derived cells lineage tracer (wpre-zsgreen) introduced in the experiment of Forte et al. These authors indeed note the existence of a myeloid-myofibroblast cluster in their scRNAseq analysis (Forte et al., 2020).

**Figure S4:**
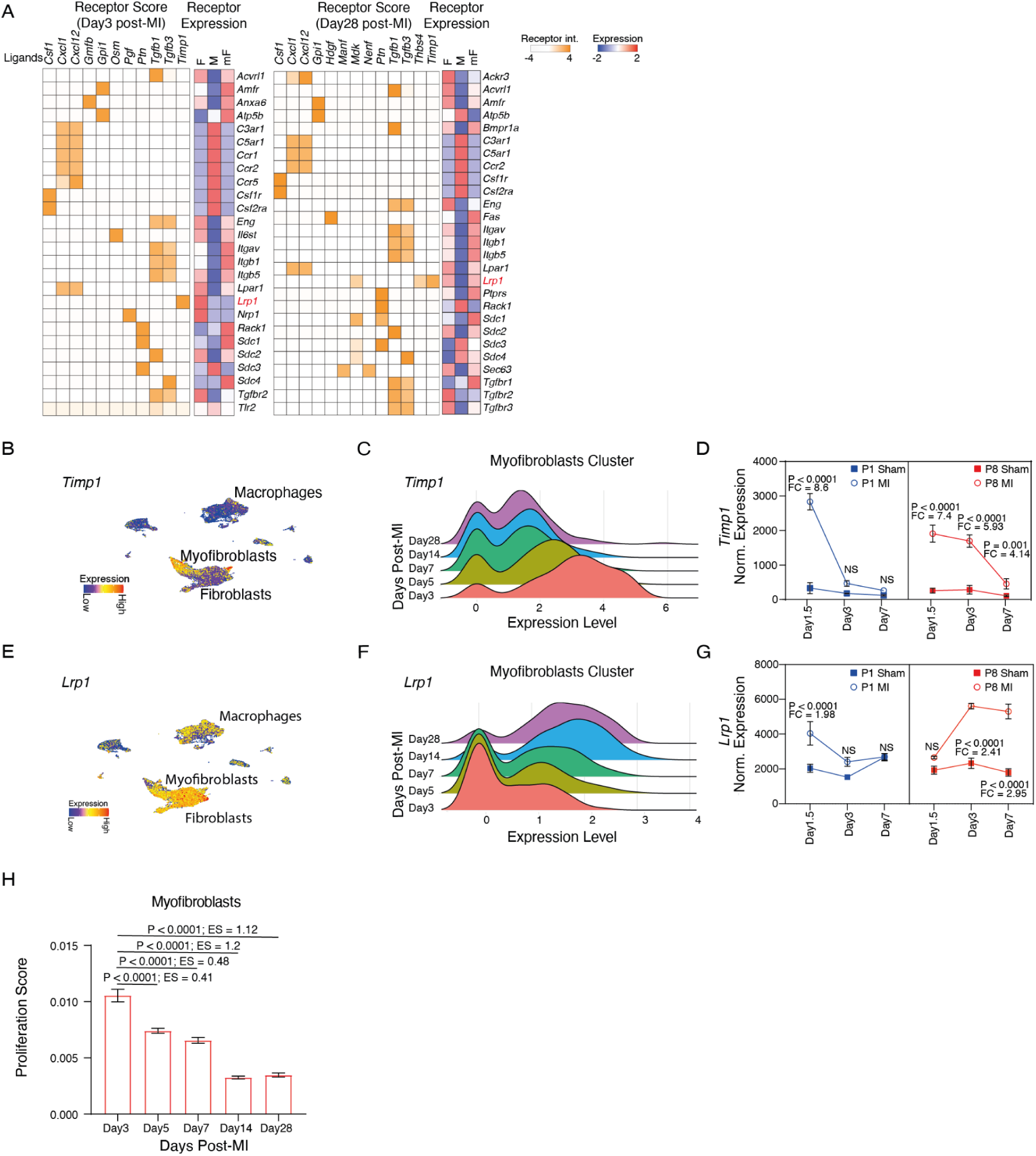
Expression of *Timp1* and its predicted receptor- *Lrp1* after MI. **(A)** Heatmap of potential receptors based on cardiac fibroblast (F), macrophage (M) and myofibroblast (mF) on days 3 (left) and 28 (right) post-MI. This data is complementary to Fig.4D. Receptor interaction score and average receptor expression per cell type are presented as mean± SD (by either white-orange scale or blue-red scale, respectively). **(B)** single-cell mRNA expression of *Timp1* by cardiac interstitial cells shows dominant expression in cardiac myofibroblasts. Scale denotes normalized mRNA expression. Dataset used is described in Fig. 1B and fig. S1A-B **(C)** *Timp1* mRNA expression dynamics after MI quantified between day 3 and day 28 in cardiac myofibroblasts cluster. **(D)** *Timp1* mRNA expression in the publicly available bulk mRNA-sequencing ((Z. Wang et al., 2019). P1 (blue) and P8 (red) mice underwent Sham (squares) or MI (circles) injury by LAD ligation and ventricles were sampled on days 1.5, 3 and 7 post-MI for sequencing. Data is shown as normalized mRNA expression. (n = 3 per time-point, per age/ treatment). Statistical analysis was performed using DESeq2 (methods), Raw P-values were adjusted for multiple testing using the procedure of Benjamini and Hochberg. **(E)** single-cell mRNA expression of *Lrp1* by cardiac interstitial cells shows dominant expression in cardiac myofibroblasts. **(F)** *Lrp1* mRNA expression dynamics after MI quantified between day 3 and day 28 in cardiac myofibroblasts cluster. **(G)** *Lrp1* mRNA expression in P1 and P8 sham and MI operated mice. **(H)** The dynamic of cardiac myofibroblasts proliferation was assessed using the scRNAseq data described in Fig. 1B and fig. S1A-B (methods). ES- effect size; FC- fold change.

